# Spatial transcriptomics defines injury-specific microenvironments in the adult mouse kidney and novel cellular interactions in regeneration and disease

**DOI:** 10.1101/2023.11.22.568217

**Authors:** Michal Polonsky, Louisa M. S. Gerhardt, Jina Yun, Kari Koppitch, Katsuya Lex Colón, Henry Amrhein, Shiwei Zheng, Guo-Cheng Yuan, Matt Thomson, Long Cai, Andrew P. McMahon

## Abstract

Kidney injury disrupts the intricate renal architecture and triggers limited regeneration, and injury-invoked inflammation and fibrosis. Deciphering molecular pathways and cellular interactions driving these processes is challenging due to the complex renal architecture. Here, we applied single cell spatial transcriptomics to examine ischemia-reperfusion injury in the mouse kidney. Spatial transcriptomics revealed injury-specific and spatially-dependent gene expression patterns in distinct cellular microenvironments within the kidney and predicted *Clcf1-Crfl1* in a molecular interplay between persistently injured proximal tubule cells and neighboring fibroblasts. Immune cell types play a critical role in organ repair. Spatial analysis revealed cellular microenvironments resembling early tertiary lymphoid structures and identified associated molecular pathways. Collectively, this study supports a focus on molecular interactions in cellular microenvironments to enhance understanding of injury, repair and disease.

**One-Sentence Summary:** Spatial transcriptomics predicted a molecular interplay amongst neighboring cell-types in the injured mammalian kidney

Main Text:

## Introduction

The mammalian kidney removes waste products from the blood, maintains fluid homeostasis and releases hormones that control blood pressure (*1*). These and other renal functions hinge on a stereotypical organization of diverse renal cell types along a cortico-medullary axis (*2*). Nephrons, the primary filtering units of the kidney, comprise over twenty distinct cell types, including species-specific sexual diversity in proximal tubule segments, that ensure recovery of key molecules from the primary glomerular filtrate (*1*, *3*). Proximal tubule cells are highly active metabolically and therefore particularly susceptible to acute kidney injury (AKI), an abrupt loss of excretory kidney function that can be caused by multiple factors including ischemia, sepsis and nephrotoxic drugs (*4*). AKI, which affects approximately 20 - 25% of hospitalized patients in the western world, is associated with increased morbidity and mortality (*5*), and development of Chronic Kidney Disease (CKD) (*6*). CKD is characterized by renal inflammation and fibrosis (*7*, *8*) and is predicted to become the fifth most common cause of death by 2040 (*9*). Despite its high prevalence and medical burden, therapeutic strategies for treating AKI and preventing the AKI-to-CKD transition are lacking.

The heightened risk of CKD following AKI has been linked to the limited regenerative capacity of the mammalian kidney (*10*). Murine models of ischemic AKI predominantly result in pronounced cell death of the proximal tubule, particularly within the S3 segment (10–12, 9, 13, 14). Injury also invokes compensatory de-differentiation and cell division of surviving proximal tubule cells, which can lead to at least partial restoration of kidney structure and function (*11*, *12*). However, even mild injury is associated with epithelial scarring, and the persistence of maladaptive epithelial cells expressing inflammation-inducing cytokines and fibrosis-associated secretory factors - inflammation, fibrosis and vascular rarefaction are key features of the pathological niche surrounding maladaptive epithelial cells (*13–24*). Several molecular and cellular features of the AKI response are conserved between mice and humans (*25–28*) and persistent pro-inflammatory and pro-fibrotic signaling are believed to promote the AKI-to-CKD progression (*29–31*).

Single cell studies have generated new insight into AKI through the identification of cell types associated with pathology and underlying transcriptional programs (*8*, *13*, *15*, *28*, *32*, *33*). However, tissue dissociation in these studies, which removes the spatial context of interacting cell types, precludes a deeper understanding of the complex cellular interplay following renal injury. In contrast, single cell resolution, spatial transcriptomics enables a readout of cell type-specific transcriptional activity while preserving information about cellular organization in tissues (*34–38*). As such, spatial transcriptomics is well-suited to provide new insight into the pathophysiological mechanisms driving progression of kidney disease and identifying cell interactions for potential therapeutic intervention.

## seqFISH provides a detailed map of the cellular composition of normal and injured kidneys and elucidates injury-specific changes

To this end, we leveraged sequential Fluorescence In Situ Hybridization (seqFISH) (*35*, *39*, *40*) which allows for the quantification of thousands of mRNA transcripts in single cells in intact tissues. seqFISH has been shown to have high capture efficiency and molecular sensitivity (*35*) and because spatial information is retained, uniquely facilitates identification of cellular subtypes and their cell-cell interactions (*35*, *41*).

To examine AKI outcomes after the initial phase of injury-invoked renal repair (*27*), we subjected C57BL/6J male mice (8-13 weeks of age) to a mild ischemia-reperfusion injury (IRI; serum creatinine levels at 48 hrs 0.3-1.7 mg/dL), waited four weeks and then collected and analyzed the kidneys. We selected 1,300 target genes for seqFISH profiling based on key cell-type enriched markers from single cell studies of normal and IRI kidney (*3*, *13*, *15*) and a long-term study of injury-associated expression during the AKI to CKD transition in mice ((*27*); Fig. 1A). Integration of control (n=3) and post AKI (n=3) samples resulted in a data set comprising 245,171 single cells, with an average of 185±153 (mean±SD) total transcripts and 87±53 individual genes detected per cell, with a strong concordance between samples (Fig. S1A-B, Fig. S2). Following rigorous quality filtering, we retained 220,753 cells for subsequent analysis. By clustering on single cell gene expression, we identified all major kidney cell types, highlighting the segmental cell diversity along the conjoined epithelial networks of the nephron and collecting system (Fig. 1B, Table S1), as well as divergent vascular endothelial sub-sets, interstitial fibroblasts and distinct immune cell types (Fig. 1B-C). The cell types identified by seqFISH were concordant with cell types previously identified using single cell RNA sequencing (scRNAseq) (Fig. S1C). Further, their spatial localization conformed to well-documented kidney anatomy (Fig. 1C-D, F, Fig. S2, Fig. S3). However, the seqFISH analysis captured a notably larger fraction of several kidney-resident cell types relative to that previously reported in scRNA-seq studies (*42*, *43*), where dissociation procedures are prone to cause populations of certain cell types to become over- or under-represented (*42*, *44*, *45*). For instance, in past work, fibroblasts were reported to comprise ∼4-10% and macrophages less than 5% in kidney single cell- and single nuclear-RNAseq datasets, whereas our data showed that fibroblasts comprised close to 20% and macrophages ∼10% of all cells (Fig. S1D). Thus, detection of the various cell types within the kidney using seqFISH is less-biased when compared to single cell-RNAseq protocols.

**Figure 1:**
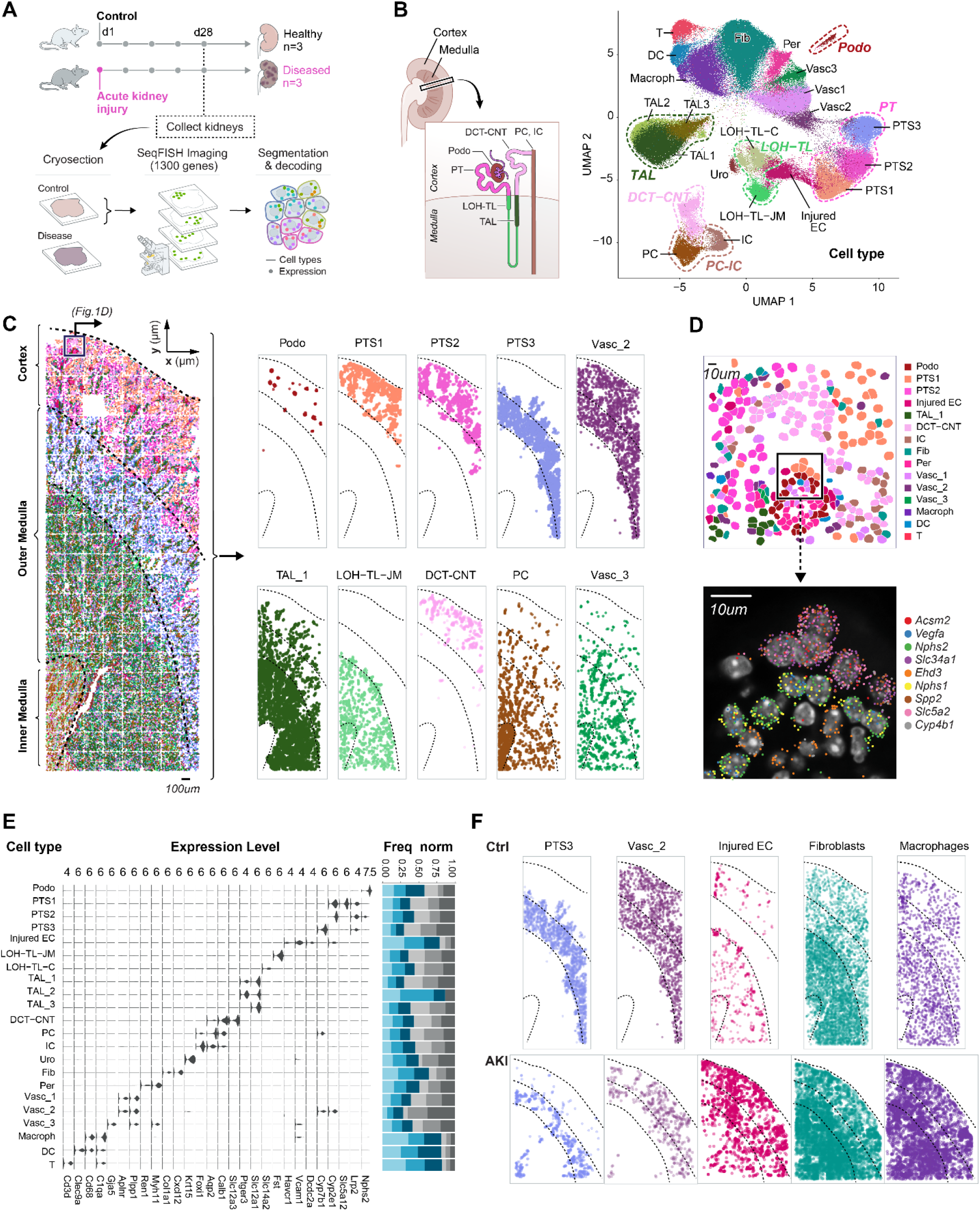
seqFISH reveals all major kidney cell types and their locations within the kidney, as well as compositional and spatial changes following AKI. A) Experimental overview: mice were subject to IRI at day 0 to induce AKI. Control and AKI kidneys were collected at day 28 and seqFISH was performed on frozen kidney sections. B) Umap depicting all cell types identified using seqFISH. Normal kidney-specific cell types are highlighted and their locations within the nephron are illustrated. DC – dendritic cells; Macroph – macrophages; DCT-CNT – distal convoluted and connecting tubule; Fib – fibroblasts; IC – intercalated cells; Injured EC – injured epithelial cells; LOH-TL-C – thin limb of the loop of Henle (comprising cells of the thin descending limb of the loop of Henle of cortical nephrons and the thin ascending limb of the loop of Henle of juxtamedullary nephrons); LOH-TL-JM – thin limb of the loop of Henle of juxtamedullary nephrons; PC – principal cells; Per – pericytes; Podo – podocytes; PT – proximal tubule; PTS1/2/3 – proximal tubule segment 1/2/3; T – T cells; TAL – thick ascending limb of the loop of Henle; Uro – urothelium; Vasc – endothelial cells. C) Spatial location of all cell types in one representative control sample. Left - all cell types, right - 10 cell types out of the total 22 that were identified when plotted individually. These cell types span the cortex and medulla, showing that seqFISH analysis captures all major cell types in different areas of the kidney. D) One representative field of view showing the cell masks color-coded by cell type (top) and a zoomed-in image showing RNA expression of representative marker genes (bottom). E) Violin plot showing the normalized expression of marker genes for each cell type. Bar plot shows the relative abundance of each cell type within the control and AKI samples. Spatial locations of five cell types within a representative control (top) and AKI (bottom) samples. PTS3 and Vasc_1 are over represented in Control, whereas Injured EC, Macroph and Fib are over represented in AKI. F) Spatial locations of PTS3, Vasc_2, Injured EC, fibroblasts and macrophages within one representative control (top) and one AKI (bottom) samples.

We next compared AKI with control samples to identify changes in the abundance and spatial locations of specific cell types. At the cortico-medullary boundary, which is known to be most sensitive to AKI (*46*), the proximal tubule segment 3 (PTS3) and vascular endothelium (Vasc_2) were strongly reduced (Fig. 1E-F; Fig. S1E, Fig. S3). Conversely, a marked increase was observed in injured proximal tubule cells, as identified by expression of *Vcam1* ((*13*, *15*); Injured EC). In addition, fibroblast, macrophage, T cell and dendritic cell populations were all elevated in AKI samples relative to control (Fig. 1E-F, Fig. S1E. Fig S3). In contrast to T cells and dendritic cells (Fig. S3), the increase in fibroblasts and macrophages was associated with a cortical expansion in injured kidney samples (Fig. 1F, Fig. S3); fibroblasts and macrophages predominated within the medulla of the uninjured kidney. In addition, we observed an over-representation of an *Slc12a1*^+^/*Ptger3*^+^ subset of thick ascending limb (TAL)-subtypes in the distal medullary loop-of-Henle (Fig. 1E; (*47*)). Thus, AKI induces global changes in both cell type composition and location.

## Distinct cellular microenvironments are specific for AKI and normal kidneys

We next sought to understand whether changes in cell populations also lead to re-organization of the local kidney architecture. Specifically, we asked whether we could detect localized microenvironments (MEs) with distinct cell type compositions within the kidney, and whether those environments changed following AKI. To this end, we calculated the frequencies of each cell type within a 30µm radius of each of the cells in the kidney tissue. We then used the resulting cell-by-neighbor matrix to cluster individual cells using Leiden clustering (Fig. 2A).

**Figure 2:**
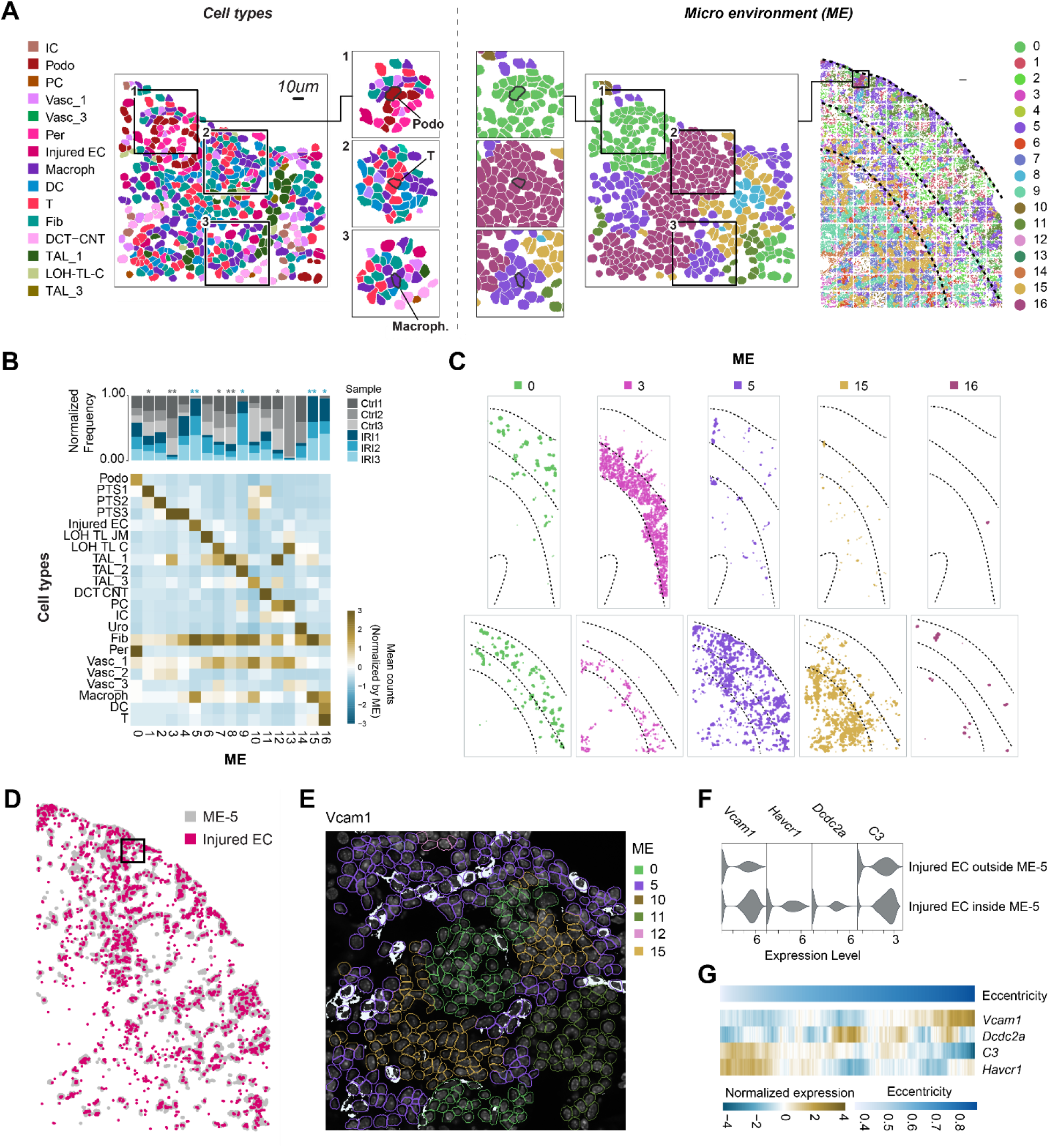
Cellular microenvironments define the spatial architecture of Control and AKI kidneys. A) Zoomed in image of one field of view in AKI sample: The cellular composition within a 30um radius around each cell is calculated and these compositions are then clustered to create 16 distinct MEs. B) Cellular composition within each ME calculated as the relative mean abundance of each cell type. Barplot showing the relative frequency of each ME within the Control and AKI samples. Gray asterisk represent enrichment in control and blue enrichment in AKI (*, p < 0.05; **, p < 0.01 of a one sided t-test). C) Spatial locations of five MEs in one representative control and one AKI sample. D) Spatial locations of all injured EC cells in one AKI sample plotted in color over the locations of all cells belonging to ME-5 plotted in gray. E) Zoomed-in image of the box plot in D showing Vcam1 antibody labeling. The cell masks are colored according to the domain assignment of each cell. F) Expression of injury marker genes in injured EC cells within ME-5 and the Injured EC cells outside ME-5 (rest). G) Normalized expression of the same genes as in F for Injured EC cells within ME-5, sorted by the eccentricity of each cell. Cells were sorted according to eccentricity values and the gene expression was averaged using a moving window with a window size of 10% total number of cells. Expression values were normalized for each gene in the heatmap.

Using this approach, we detected 17 MEs with a distinct cell type composition and spatial location (Fig. 2B-C, Fig. S4A, Fig S5). Eight MEs were not markedly different between individual control and AKI samples, five were enriched in the control kidneys and four were specific to the AKI samples (Fig. 2B, Fig. S5). To limit sampling bias, we focused our analysis on MEs that were equally enriched or depleted in all control and AKI replicates (indicated by asterisk in Fig. 2B, gray – enriched in control and blue – enriched in AKI). In Figure 2C, we show examples of MEs which are similar between control and AKI (ME-0), absent from AKI (ME-3) or enriched in AKI (ME-5, ME-15, ME-16). AKI-specific MEs were largely a combination of injured epithelium, fibroblasts and immune cells in varying proportions, and were mostly depleted of the primary tissue cells. Interestingly, dendritic cells, macrophages, T cells and fibroblasts were distributed across different MEs in the control samples, but concentrated within specific MEs in AKI samples (Fig. S4B). Therefore, the emergence of these cells, whether by differentiation, proliferation or recruitment from the blood, results in redistribution into distinct environments upon AKI.

## Characterization of the pathogenic niche in AKI

Recent studies in mouse and human have drawn attention to the tissue micro-environment around injured proximal tubule cells, which have been associated with renal pathology (*13*, *15*, *28*). Injured proximal tubule cells display the cell adhesion molecule Vcam1 and adopt a senescence-associated secretory phenotype (SASP) (*13*, *15*, *28*, *33*). We identified a similar population of *Vcam1*-positive epithelial cells in our data set by unsupervised clustering (Cluster Injured EC, Fig. 1B, E). The majority of these injured epithelial cells reside within a single ME (ME-5, Fig. 2B, D). As a means of internal validation, we stained with anti-Vcam1 antibody concurrently with seqFISH analysis showing Vcam1 protein is mainly present in Injured ECs within ME-5 (Fig. 2E). In addition, the Injured ECs within ME-5 were associated with stronger expression of several other injury-associated genes when compared to Injured ECs outside of ME-5 (Fig. 2F).

Proximal tubule cells undergo a de-differentiation in response to AKI that manifests in several morphological changes including a loss of the apical brush border and flattening of the epithelium (*48*). To assess the nuclear morphology of Injured ECs within ME-5 and correlate any changes with expression of injury genes, we estimated the eccentricity of Injured EC cell masks, in which an eccentricity value of 0 represents a perfect circle and values between 0 and 1 represent an ellipse. We found that eccentricity analysis could sub-divide Injured EC cell expression profiles. With increasing eccentricity values (corresponding to a flattening of the nucleus), *Vcam1* expression increased, while expression of other injury-related genes did not correlate with high eccentricity (Fig. 2G). Taken together, our findings show that ME-5 represents an injured and likely pathogenic niche. The majority of injured ECs, and injury associated fibroblasts, vasculature and immune cells, reside within this niche, and morphological change within the injured epithelium associated with elevated expression of injury-response genes.

## *Clcf1 - Crlf1* interactions between injured EC and fibroblasts shape the injured niche

We sought to identify signaling between Injured EC and the other cell types present within ME-5 (fibroblasts, macrophages, T cells, DCs, Vasc_1, few normal PT cells), which are potential drivers of the organization of the injured niche. We first identified predicted signaling initiated by the Injured ECs using Nichenet (*49*) and found that *Clcf1* – *Crlf1* signaling was upregulated between Injured ECs and fibroblasts within the AKI samples (Fig 3A, Fig S6A). Upregulation of *Crlf1* and *Clcf1* in kidney disease has been indicated (*50–52*), and *Crlf1* was shown to be upregulated in the fibrotic lung (*53*). *Clcf1* encodes a member of the Il-6 cytokine family which is thought to engage *Crlf1* as a chaperone in signaling through the receptor *Cntfr* (*54*, *55*). While *Clcf1* was specifically expressed within injured ECs (Fig 3A, Fig S6A, Fig. S7), *Crlf1* was predominantly expressed in a distinct subset of ME-5 associated fibroblasts, not previously identified in sc/snRNA-seq studies (Fig 3A-B, Fig. S6B, Fig. S8).

**Figure 3:**
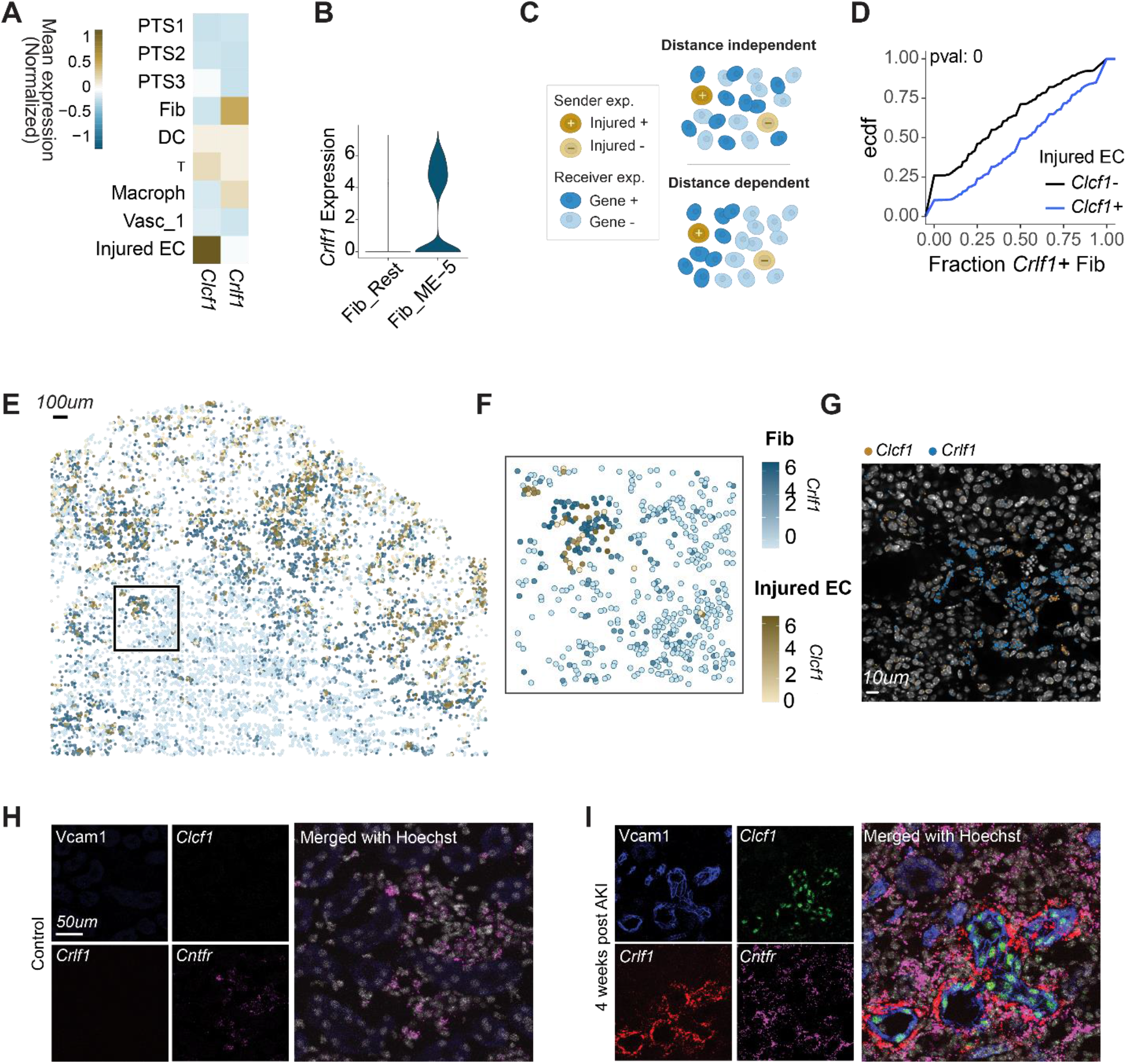
Injured EC are interacting with macrophages and fibroblasts within ME-5. A) Mean expression of *Clcf1* and *Crlf1* on the cell types present in ME-5 B) *Crlf1* expression on fibroblasts within ME-5 (Fib_ME-5) and outside ME-5 (Fib_Rest) C) Illustration showing cellular behaviors under distance-dependent and distance- independent signals. D) ECDF plots of the fraction of *Crlf1*+ fibroblasts within a 30μm radius around *Clcf1*+ or *Clcf1*-Injured EC. p-value was calculated using a one-sided student’s t-test. E) Spatial expression of *Clcf1* on Injured EC (gold) and its receptor *Crlf1* (blue) on fibroblasts. F) Zoomed-in image on the inset in C showing *Clcf1* on Injured EC and *Crlf1* on fibroblasts. G) Zoomed-in image showing mRNA dots of the corresponding genes. H) Antibody-staining of the Injured EC marker Vcam1 coupled with RNAscope evaluation of *Clcf1*, *Crlf1* and *Cntfr*, the receptor for the *Clcf1-Crfl1* complex. Staining was done on a control sample. I) Same as in H, measured on AKI sample.

Our analysis also identified upregulation of *Cxcr6* in T cells and *Itgax* in DCs within AKI samples suggestive of Injured EC signaling to T cells and to dendritic cells through *Cxcl16* - *Cxcr6* and *Icam1* – *Itgax* signaling axes, respectively, recruiting immune cells within AKI (Fig. S6A). In the context of liver fibrosis, *Cxcr6*-dependent recruitment of CD8 T cells and NKT cells is known to contribute to disease progression (*56–58*) and a similar role has been suggested for a *Cxcl16* - *Cxcr6* axis in kidney fibrosis (*59–61*). Macrophages in AKI also express *Cxcl16,* consistent with a contribution to T-cell recruitment (Fig. S6A). *Csf1 - Csf1r* signaling axis between injured ECs and macrophages was also upregulated within AKI samples (Fig. S6A), in line with previous studies that link injured epithelial *Csf1* production to macrophage-mediated recovery following AKI (*62*). Analysis of two published scRNAseq and snRNAseq datasets of dissociated human and mouse kidney samples showed that the elevated expression of the top ligands identified here including *Clcf1* and *Csf1* as well as *Cxcl16*, *Icam1* and *Tgfb2* was conserved across species (Fig. S6B).

We hypothesized that spatially localized ligand-receptor interactions (spatially dependent; Fig. 3C) will contribute specifically to the structuring of the microenvironment around and via injured ECs as opposed to signaling with no spatial preference (spatially independent; Fig 3C). To assess the spatial localization of the signals, we quantified the fraction of receptor-expressing target cells within a 30um radius of Injured ECs with or without its cognate ligand expression, regardless of ME assignment. Figure 3D shows the Empirical Cumulative Distribution Function (ECDF) of the fraction of *Crlf1*+ fibroblasts within the vicinity of *Clcf1*+ injured ECs compared to *Clcf1*− injured ECs. This analysis detected a strong spatial preference for *Crlf1*+ fibroblasts to *Clcf1+* injured cells (p-value of a one-sided students t-test = 0, gene expression in one representative AKI sample is shown in Figure 3E-G). Similarly, no, or only a small, spatial preference was detected for the other ligand-receptor pairs involved in signaling from Injured ECs to fibroblasts, vasculature and different immune cell types (Fig. S6C). To validate the predicted expression and spatial distribution of the *Clcf1*-*Crlf1* pairings, we combined anti-Vcam1 immunodetection with serial RNA-FISH on control and AKI samples. Consistent with seqFISH analysis, *Crlf1*+ cells were specifically concentrated adjacent to *Clcf1*+Vcam1+ Injured ECs (Fig. 3H-I, Fig. S9). Contrary to the specific localization of *Crlf1* around injured cells, *Acta2,* a well-documented marker of inflammatory myofibroblasts (*15*), exhibited heterogeneous expression (Fig. S10). *Clcf1*, the predicted target of *Crfl1* activity, signals through *Cntfr*. *Cntfr* expression colocalized with *Crlf1* in peri-epithelial myo-fibroblasts but was not limited to this population (Fig. 3H-I).

Taken together, our findings implicate injured EC signals in remodeling of vasculature, recruitment of immune cells and reshaping of the fibroblast population within the AKI samples. Using our spatial data, we show that while several signaling events are upregulated within AKI samples, only *Clcf1-Crlf1* interactions are highly spatially localized. Signaling which is upregulated but does not show spatial localization could present global changes following AKI or past events leading to the formation of the injured niche. Our combined spatial data and validation suggest that *Crlf1* is a specific identifier of fibrotic processes closely coupled to Injured ECs, and that *Clcf1 - Clcf1* interaction with fibroblast cells are a constitutive defining feature of the injured niche.

## Fibroblasts show distinct expression patterns between different MEs

To determine whether gene expression changes could be identified more precisely in different MEs, we analyzed fibroblasts within the four AKI-specific MEs (ME-5, ME-9, ME-15 and ME-16). Both composition of cell types and location in the tissue were distinct for each ME (left panel in Fig. 4A). ME-5 localized to the cortex and corticomedullary boundary, ME-15 extended from the corticomedullary boundary to the medulla, ME-9 was concentrated around TAL_2 cells in the medulla and ME-16, comprising predominantly immune cell types, was scattered throughout the kidney (Fig. 4A, Fig. S5).

**Figure 4:**
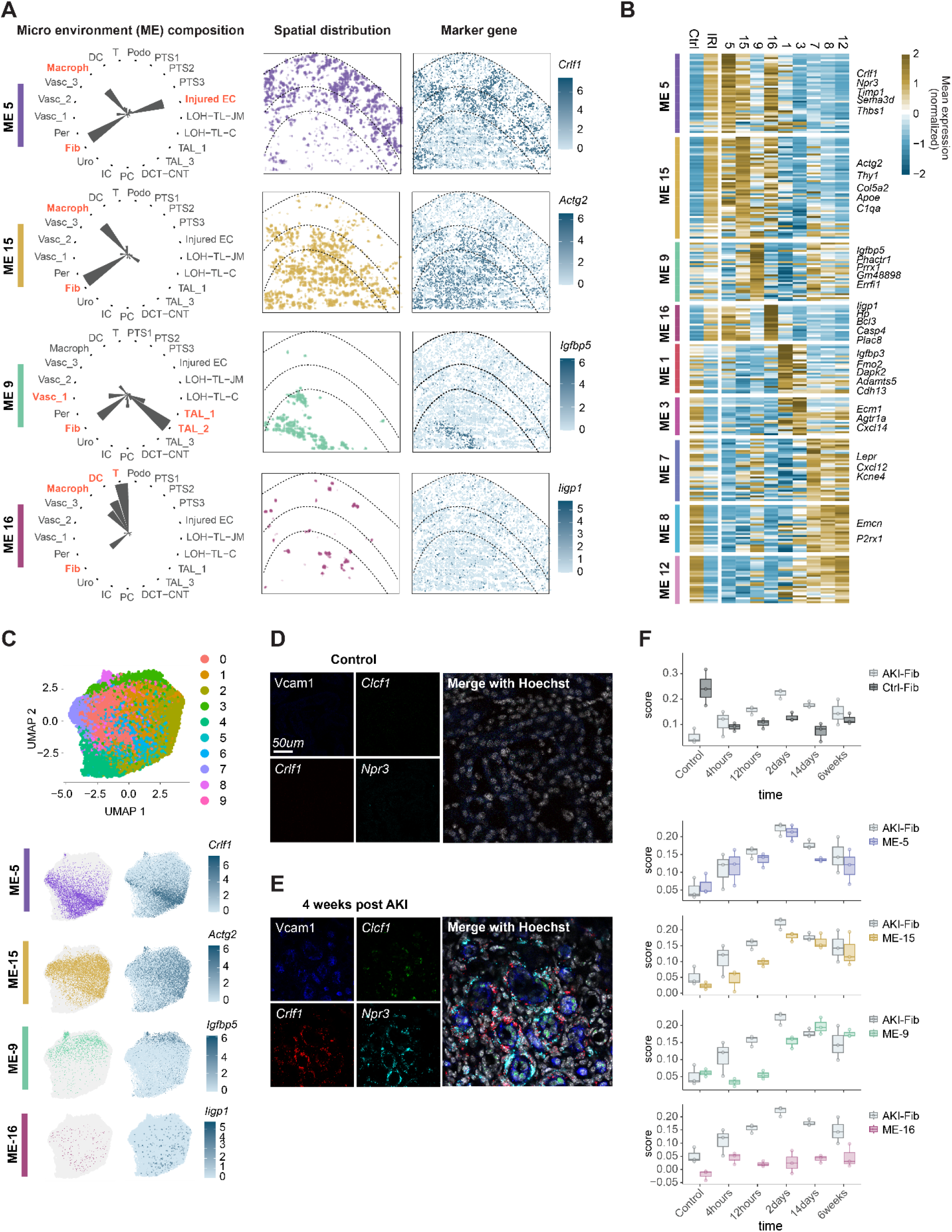
Within different MEs, Fibroblasts have distinct gene expression patterns, with a subset of Fibroblasts representing acute response to injury. A) Left - coordinates plot showing the composition of four MEs enriched in AKI samples. Middle - spatial locations of each of the MEs in one AKI sample. Left - expression of marker genes of each ME in all Fib in the sample. B) Mean expression of the ME differentially expressed genes over each MEs as well as control and AKI. The depicted MEs represent four enriched in AKI (ME-5,15,9,16) and five enriched in control (ME-1,3,7,8,12). C) Top – Leiden clusters only of Fibroblasts based on gene expression presented in Umap space. Bottom – same ME and marker genes as in A, presented on Umap space. D) Vcam1 antibody-staining with *Clcf1*, *Crlf1* and *Npr3* expression measured using RNAscope on control sample. E) Same as in E on AKI sample F) Average expression score for the differentially expressed genes shown in B in a time course snRNAseq data. The score was calculated for single cells using Seurat and averaged for each mouse within the snRNAseq time course experiment. Dots represent individual mice.

The top enriched fibroblast gene-sets for each ME, shown in Figure 4A-B, revealed distinct ME-associated gene expression (see Table S1 for full list of differentially expressed genes). Figure 4A shows that the expression of the top ME-associated genes is highly restricted to the location of each ME in the physical space. Interestingly, we found that many of these genes were not detected when clustering fibroblasts based on gene expression alone without considering spatial ME information (Figure 4C-top). Out of the four ME-specific fibroblast maker genes *Crlf1*, *Actg2*, *Igfbp5* and *Iigp1*, only *Igfbp5* was enriched in the gene expression-based cluster (cluster 3 in Figure 4C-top), while the other genes did not show cluster-specific expression. These data underscore the importance of considering the spatial environment of fibroblasts to identifying genes with a potential functional significance.

In addition to the expected fibroblast markers, we found that *Crlf1*, *Npr3* (which encodes the natriuretic peptide receptor 3) and *Timp1* (which encodes TIMP metallopeptidase inhibitor 1) were markedly enriched in the ME-5 fibroblasts (Fig. 4A-B). In previous work, *Npr3,* a known blood pressure regulator, reduced disease severity upon inhibition in a model of cardiac fibrosis (*63*), whereas upregulation of *Timp1* was reported to increase kidney scarring (*64*, *65*). Here, we validated co-expression of *Crlf1* and *Npr3* in association with injured ECs using immunostaining and serial RNA-FISH (Fig. 4D-E). In ME-15, which comprises fibroblasts and macrophages, *Actg2*, encoding a member of the smooth muscle actin family, was the top gene identifier of ME-15 fibroblasts (Fig. 4A). *Actg2* has not previously been associated with kidney disease, although a recent study identified *Actg2* as a candidate in long term kidney impairment following acute decompensated heart failure (*66*). ME-15 fibroblasts also elevate expression of *Apoe* and *C1qa,* key inflammatory genes highly expressed by macrophages, suggesting both macrophages and fibroblasts contribute to the inflammatory environment. Fibroblasts in ME-16 were distinguished by expression of *Ligp1,* which encodes the interferon-induced GTPase1, and expression of *Bcl3,* an inhibitor of apoptosis, and the caspase *Casp4*. In contrast to ME-5, −15 and −16 fibroblasts, ME-9 fibroblast marker genes were not highly expressed within the total AKI fibroblast population and several genes were higher in control fibroblasts. One of the most strongly enriched fibroblast genes in this ME was the insulin growth factor binding protein (IGFBP), encoded by the *Igfbp5* gene. IGFBPs have been linked to CKD progression (61) and disease progression in a mouse model of diabetic kidney disease (*67*).

## Marker genes for fibroblasts within the MEs show distinct temporal expression in sn-RNAseq data

Given the highly distinct gene signatures of fibroblasts across the different MEs, we next asked if expression of the identified ME-specific fibroblast marker genes changes along the AKI-to-CKD transition. To answer this question, we analyzed published snRNA-seq data collected at six time points post AKI (2 hours, 4 hours, 12 hours, 2 days, 14 days and 6 weeks; (*15*)), extracted the fibroblast cluster from this data set and calculated an ME score for each cell (combined expression score of all fibroblast marker genes for each of the AKI-specific MEs). Figure 4F depicts the average ME score for fibroblasts in each mouse at each time point with dots representing individual mice. The top plot represents the average scores for fibroblast marker genes in control and AKI samples, regardless of ME assignment.

As expected, following AKI, the control-fibroblast score is reduced immediately and remains relatively low until 6 weeks post AKI, while the AKI-fibroblast score is low in control samples, but increases and peaks at day 2 after AKI, remaining elevated at 6 weeks. This timeframe suggests that the injury induces fibroblast differentiation and recruitment, as captured in the AKI-fibroblast score, and that AKI-induced fibroblasts persist 6 weeks after injury.

We found divergent patterns in the ME-specific fibroblast scores (Fig. 4F). ME-5 and ME-15 scores were similar to the AKI-fibroblast score, peaking at day 2 post AKI, with ME-15 showing lower score values than ME-5 whereas the ME-9 score was significantly delayed and peaked at 14 days post AKI, suggesting that this fibroblast subset appears late following AKI. Although ME-16 genes were highly specific to AKI samples in our data (Fig. 4B), the ME-16 score was low throughout the time course, potentially reflecting detection issues due to sampling bias or population size in the snRNA-seq data set. Assuming that the snRNA-seq data is capturing cells from different sites of the tissue, our analysis suggests that the abundance of different fibroblast subsets detected reflect changes along the AKI-to-CKD transition. Thus, beyond spatial mapping, seqFISH analysis provides additional molecular granularity as to how cell populations acquire distinct properties during a biological process.

## Immune cells show distinct spatial distributions which correlate with phenotype and inflammatory potential

Next, we focused on immune cells as drivers of inflammatory fibrosis. To identify T cell subtypes present in our data, we mapped T cell populations onto a reference data set of mouse T cells (*68*), using SEURAT. We identified known T cell populations including CD4 and Tregs (high *Cd83* and *Ctla4* expression), naïve, effector (high *Cxcr3* expression and expression of cytotoxic genes such as *Nkg7)*, and exhausted (no Cxcr3 expression but expression of the inhibitory molecule *Pdcd1*) CD8 T cells (Fig. 5A). When we quantified the fraction of each T cell subtype within MEs −5, −15 and −16 in the AKI samples, we found a difference in the fraction of CD4 T cells amongst all T-cells: ∼40% in ME-5, 50% in ME-15 and 55% in ME-16 (Fig. 5B). We noticed a similar trend for Tregs, although, as expected, Tregs represented a smaller subset of the T cell population (Fig. 5B). Strikingly, the inverse trend was apparent for effector CD8 T cells, while no clear trend was evident for naïve and exhausted CD8 cells (Fig. 5B).

**Figure 5:**
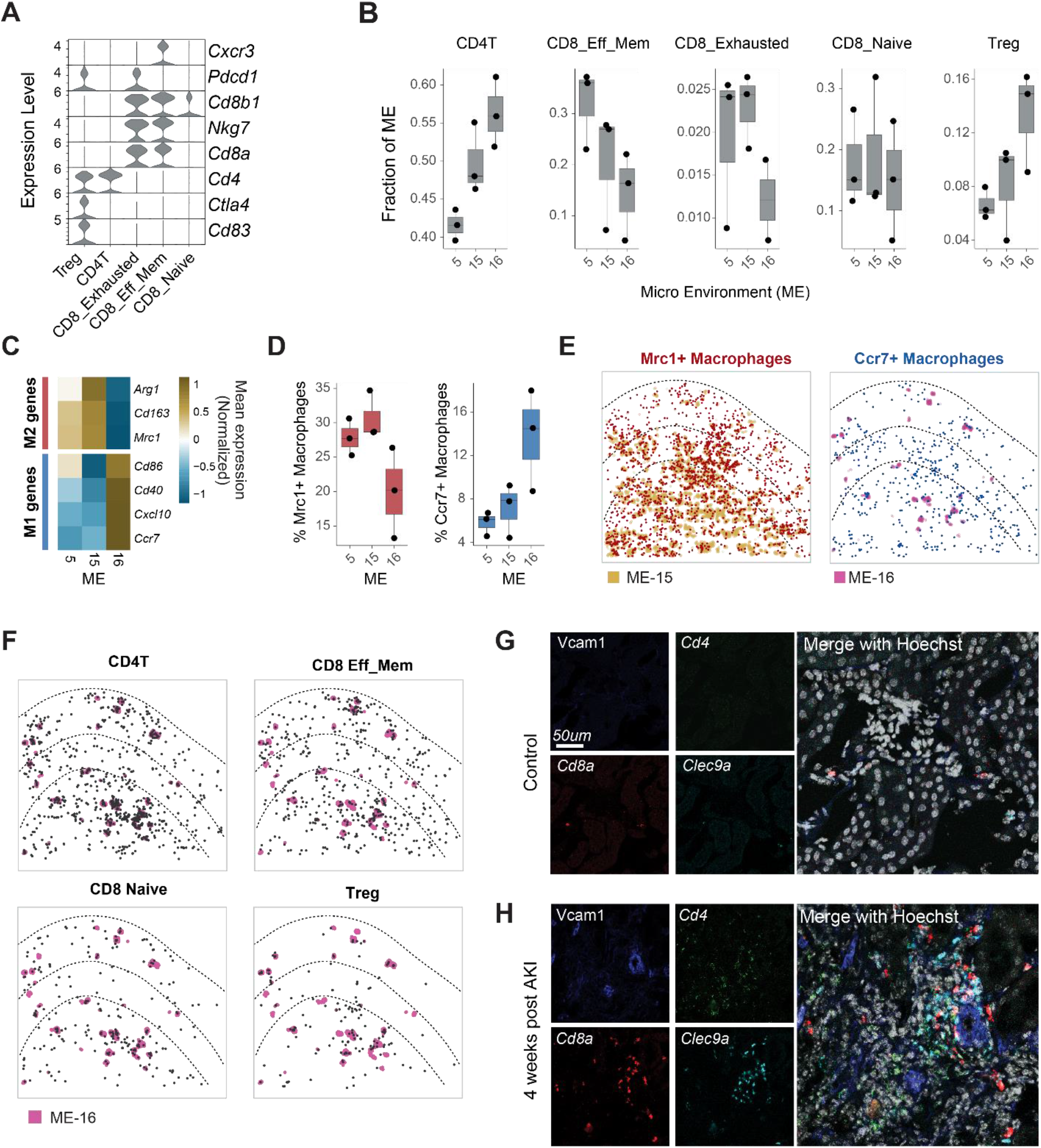
T cell subtypes preferentially reside into different MEs, correlating with their functionality. A) All T cells in our dataset were mapped to a tumor-infiltrating lymphocyte dataset. The T cells were mapped to six known cell populations - helper T cells (CD4T), cytotoxic T cells (CD8T Eff_mem), naive and exhausted CD8 (CD8_Naive, CD8_exhauseted) and regulatory T cells (Treg). The violin plot shows marker genes expression for each of the T cell subtypes in our data. B) Fraction of T cells of each subtype in IRI enriched MEs (5, 15, 16) for the three AKI mice. Each dot represents one mouse. The fraction is the total fraction of each subtype out of all T cells within the specified domain. C) Average expression of three M1 (blue) and four M2(red) markers within Macrophages in the AKI enriched MEs. D) Fractions of Mrc+ (left) and Ccr7+(right) Macrophages within the AKI enriched MEs boxplots showing distribution of mean values for the three AKI samples. Dots represent individual mice. E) Spatial distribution of the same genes as in D. Left - ME-15 is indicated and Mrc+ cells are colored in red. Right - ME-16 is indicated and Ccr7+ cells are colored in blue. F) Spatial locations of the T cell subtypes within one AKI sample. ME-16 is depicted in color. G) Vcam1 antibody staining as well as the markers *Cd4*, *Cd8* and the DC marker *Clec9a* are detected with RNAscope on a control sample. H) Same as in G for an AKI sample

Since macrophages represent a large fraction of the immune cells in AKI-specific MEs, we measured marker genes linked to differential activation states of M1 and M2 macrophages. In general, M1 macrophages promote disruptive, disease-associated inflammatory responses whereas M2 macrophages promote constructive, inflammatory-associated tissue repair (*69*). We found that the average expression of the M2-related genes *Mrc1*, *Cd163* and *Arg1* was increased in macrophages within the ME-5 and ME-15 groupings, while M1-related genes *Cxcl10, Ccr7, Cd40* and *Cd86*, were expressed at higher levels on the average in ME-16 macrophages (Fig. 5C). In agreement with these data, *Mrc1*+ macrophages were more prevalent in ME-5 and ME15, and *Ccr7*+ macrophages were more prevalent in ME-16 (Fig. 5D-E).

Tertiary Lymphoid Structures (TLS), increasingly recognized as contributors to chronic inflammation (*22*), are organized lymphoid aggregates that form ectopically in response to a disturbance in tissue homeostasis and then act as localized hubs that enables signal exchange between immune cells, which in turn promotes development of adaptive immunity within a tissue. Unlike secondary lymphoid structures (such as the lymph nodes and the spleen), TLS are unencapsulated, exposing cells within to multiple signals from the environment. In the context of autoimmune and chronic inflammation, the presence of TLS is correlated with severe disease, and in the context of chronic kidney disease, with more severe inflammation and fibrosis (*20*). Mechanistically, the emergence of TLS in a long-term model of the AKI to CKD transition was linked to maturation of B cell-directed autoimmunity against the target kidney tissue (*27*, *70*).

Interestingly, the cellular composition and appearance of ME-16 shows characteristics of a nascent TLS. ME-16 is composed of T cells, DCs and macrophages with a small number of fibroblasts (Fig. 4A). ME-16 clusters also appear scattered without a preference for the cortex or the medulla (Fig. 2C and Fig. 4A). When overlaid on the locations of ME-16 (Fig. 4F), the spatial distribution of CD4 and CD8 T cells were dispersed within and outside of ME-16, with CD4 T cells clearly aggregated in ME-16. Thus, ME-16 has expected properties of an early TLS in which CD4 T cells and Tregs are providing instructive signals to CD8 and other immune cells. Further, employing RNA-FISH detection of *Cd4*, *Cd8a* and the DC marker *Clec9*, we detect early aggregation of DCs, CD4 and CD8 T cells 4 weeks following AKI (Fig. 5G-H and Fig. S11). CD8 T cells trend towards cortical prevalence and CD4 towards medullary enrichment. Moreover, elevated *Cxcr6* expression in CD8 T cells (Fig. S12) suggests interaction with *Cxcl16*-expressing Injured ECs (Fig. S6A).

Although the clear enrichment of T cells and DCs within ME-16 is consistent with localized antigen presentation, we did not detect any B cells, in line with kidney profiling data showing a later engagement of the B cell response (26, 70). Because M1 macrophages are preferentially enriched within ME-16, we propose that ME-16 represents an early lymphoid structure that propagates inflammation. In contrast, the presence of M2 macrophages in ME-15 and ME-5 suggests that these MEs are undergoing a combination of inflammatory and fibrotic processes where the inflammatory response is sequestered and replaced by fibrotic processes (*71*).

## Discussion

Cell identity is largely defined by gene expression profiles. However, unique microenvironmental interactions within tissues also represent important determinants of both gene expression and cell identity. To gain a more granular understanding of how cell identity and function is shaped by such micro-level interactions, we need to incorporate information about gene expression with cellular locations within tissues. Transcriptionally heterogeneous cellular populations characterized by scRNA-seq analysis are typically clustered into sub-populations. However, without a specific reference to a corresponding purified cellular subtype, this approach is prone to over- or under-clustering, and may even fail to identify important cellular populations (*41*). Spatial transcriptomics overcomes this limitation as cells are first clustered globally into well-characterized cell types, and this information is then integrated with the cellular locations to identify cellular microenvironments (MEs) for each individual cell. Thus, each cell now has two identifiers – cell-type, based on its gene expression, and a cellular ME, based on the composition of its close neighbors.

We leveraged seqFISH to study cellular and structural changes following kidney injury, which dramatically disrupts the highly complex kidney architecture and triggers two parallel responses: regeneration of the damaged epithelia and fibrosis leading to development of CKD. Based on our analysis, we have created a comprehensive map of the cellular, molecular and structural changes following AKI, which can be leveraged to enhance our understanding of pathophysiologic processes underlying the AKI-to-CKD transition. To ease data accessibility for the scientific community, we have created an interactive website through which the data can be explored (https://woldlab.caltech.edu/celltiles/mouse_kidney_fibrosis/).

Our approach allowed us to gain a deeper understanding of an injured ME undergoing responses specific for the AKI-insult; here referred to as ME-5. Within ME-5, we identified injured ECs, fibroblasts, macrophages as well as immune cells (T, DC) and normal kidney cell types (PTS1, PTS2, PTS3 and vasculature). Injured cells within the ME expressed specific ligands, which were conserved in human injured ECs. When we used spatial coordinates to identify specific ligand-receptor interactions that were highly spatially localized, we found that *Clcf1-Crlf1* interactions were upregulated within the injured ME, and spatially localized between injured ECs and fibroblasts. This finding suggests that *Crlf1* expression within fibroblasts is a highly specific determinant for the fibrotic processes surrounding the injured cells. At a broader level, our approach illustrates that analyzing localized interactions at a microenvironmental level can help us gain a multi-faceted view of how cell-cell communication contributes to tissue organization under both normal and injury settings.

Following injury, global gene expression and cellular populations differed across the emerging MEs. Importantly, we detected gene markers delineating ME-specific fibroblasts subpopulations that we did not detect by our initial gene expression-based clustering. Specifically, we found that *Crlf1* together with *Timp1* and *Npr3* are specific markers for fibroblasts localized around injured cells, possibly contributing to fibrosis. Conversely, *Actg2*, *Apoe* and *C1q* labeled fibroblasts which were more medullary and distant from the injured cells, suggesting that these propagate inflammation. Interestingly, we identified ME-16, a specific microenvironment which resembled a tertiary lymphoid structure both in terms of morphology and cell type composition. In this ME, we found preferentially CD4 T cells and Tregs, as well as macrophages with a pro-inflammatory gene expression. Thus, our spatial analysis was able to identify discrete cellular subsets not identified by gene expression alone and revealed a morphological structure with potential functional importance.

Taken together, we here leveraged spatial information to identify localized inter-cellular interactions and fibroblasts subtypes with potential functional roles in injury progression that were not identifiable by gene expression data alone. We envision that this kind of data can be leveraged beyond constructing spatial domains where correlations between cell types and gene expression within their neighbors can be used to find continuous changes and flow of information between cells within the tissues.

## Limitation of this study

Although our extensive gene panel allowed us to detect all major kidney cell types as well as location-specific gene expression patterns and intercellular interactions, our panel is still limited. To comprehensively capture the consequences of intercellular interactions there is a need to identify additional molecular pathways and therefore scale up the number of measured genes in future studies. In addition, since intercellular signaling was inferred from RNA expression of ligands and receptors, identified molecular interactions will have to be validated using additional approaches in future studies. We have demonstrated that the identification of MEs captures the kidney morphology and is highly coherent between the different samples. However, similarly to cell type clustering, the process of ME identification required thresholding the data (30um neighborhoods) and manual curation, which could potentially introduce biases.

Our analysis suggests that, upon injury, specific fibroblasts populations appear in earlier timepoints and TLS-like MEs, which can develop into larger TLS, form in later timepoints. To verify these hypotheses, performing a time-course experiment ranging from very early timepoints following injury (2 days) and up to several months would be required. This can be addressed in future work.

## Supporting information

DEG cell types, DEG fibroblasts between MEs

Codebook and probe info for cell markers

Codebook and probe info for genes

Final serial probes

## Acknowledgements

We thank Inna-Marie Strazhnik for help with figure illustration and design.

## Author contributions

MP, LMSG, APM and LC conceptualized the study. JY and MP designed the probes with input from LMSG. LMSG and KK performed surgeries, tissue collection and validation experiments. JY performed seqFISH experiments. KLC and MP wrote image processing scripts. MP, LMSG and SZ analyzed data with input from GC MT, APM and LC. MP, LMSG and HL were responsible for visualization. MP and LMSG wrote the initial draft of the manuscript. MP, LMSG, APM and LC edited and reviewed the manuscript. APM and LC supervised all aspects of the project.

## Data and materials availability

Scripts used for pre-processing seqFISH images can be found at https://github.com/CaiGroup/pyfish_tools. The source data and processed data from this study will be available https://datadryad.org/stash after completion of the review process. The codebook is available in Table S3. Processed data can be browsed at https://woldlab.caltech.edu/celltiles/mouse_kidney_fibrosis/.

## Supplementary Materials

### Materials and Methods

#### Readout probe design and synthesis

Readout probes 15 nt in length were designed as previously described (*72*). In brief, a set of probe sequences was randomly generated with combinations of A, T, G or C nucleotides. Readout-probe sequences within a GC-content range of 40–60% were selected. We performed a BLAST search against the mouse transcriptome to ensure the specificity of the readout probes. To minimize cross-hybridization of the readout probes, any probes with ten contiguously matching sequences between readout probes were removed. The reverse complements of these readout-probe sequences were included in the primary probes according to the designed barcodes. The fluorophore-coupled 15-nt readout probes (Alexa 488, 647 (Thermo Fisher Scientific) and Cy3B (GE Healthcare).

#### Primary probe design

Primary probes were designed as previously described (*35*, *73*) with some modifications. Probe length was set to 35nt. Any gene with less than 16 probes was discarded. Housekeeping genes and highly expressed genes were discarded to avoid optical crowding of the fluorescence probes. The probes were divided into two pool designs: the first pool comprised 131 cell marker genes; this pool was barcoded with 4 pseudocolors to be decoded over 16 hybridization rounds in total. The second pool contained the rest of the genes and was barcoded with 9 pseudocolors to be decoded over 36 hybridization rounds in total. Each pool was read in three channels, however we used the seqFISH+(*35*) design scheme where a set of genes was only decoded over one channel to avoid chromatic aberration. The first cell marker pool was designed such that there were 44, 43 and 44 genes barcoded in the first, second and third fluorescent channels respectively. The second pool contained 390, 398 and 390 genes in the first, second and third channels respectively.

#### Serial probe design and hybridization

To ensure detection of the injury specific markers Vcam1 and Havcr1, we created primary probes which were not barcoded for these two genes. Vcam1 was targeted with 30 probes and Havcr1 with 27 probes. The probes were designed such that each one carried two binding sites for the same readout (without splitting into 4 readout barcode). The probes were obtained from IDT and no further synthesis was performed. Probes were hybridized on the samples together with the barcoded probe pool at a concentration of 10nM per probe.

#### Primary Probe synthesis

Primary probes were generated from oligoarray pools (Twist Bioscience) as previously described (*35*). In brief, probe sequences were amplified from the oligonucleotide pools with limited two-step PCR cycles and PCR products were purified using QIAquick PCR Purification Kit (Qiagen 28104). Then, *in vitro* transcription (NEB E2040S) was performed followed by reverse transcription (Thermo Fisher EP0751). After reverse transcription, the single-stranded DNA (ssDNA) probes were alkaline hydrolyzed with 1 M NaOH at 65 °C for 15 min to degrade the RNA templates and neutralized with 1 M acetic acid. Finally, probes were ethanol precipitated, and eluted in nuclease-free water.

#### Coverslip functionalization

Coverslips were cleaned with a plasma cleaner on a high setting (PDC-001, Harrick Plasma) for 5 min. Then coverslips were rinsed with 100% ethanol three times, and heat-dried in an oven at >90 °C for 30 min. Next, the coverslips were treated with 100 μg μl^−1^ of poly-D-lysine (P6407; Sigma) in water for >3 hrs at room temperature, followed by three rinses with water. The coverslips were then air-dried and kept at 4 °C for no longer than 2 weeks.

#### Mice

Mouse handling, husbandry and surgical procedures were performed according to the guidelines of the Institutional Animal Care and Use Committee at the University of Southern California (protocol numbers 11911).

#### Ischemia-Reperfusion Injury model and tissue collection

Warm bilateral renal ischemia-reperfusion injury (IRI) was performed as previously described on male C57BL/6J (weight 26-27g, age 8-13 weeks) (*27*). Male non-surgery C57BL/6J mice were used as controls. Kidneys were collected 28 days post IRI. After organ perfusion with ice-cold RNase-free phosphate buffered saline (PBS), the kidney capsule was removed and the kidneys were fixed in RNase-free 4% paraformaldehyde-PBS overnight at 4°C. Kidneys were then equilibrated overnight in RNase-free 30% sucrose-PBS, embedded in O.C.T. compound (Tissue-Tek) in a dry-ice ethanol bath and stored at −80°C. Ten micrometer cryostat sections of kidneys were placed on the functionalized coverslips on dry-ice for subsequent seqFISH experiments. Secondary validation experiments used sections adjacent to those analyzed by seqFISH.

#### Probe hybridization

Tissue sections were permeabilized in 70% ethanol at −20 °C for >1 h, cleared with 8% SDS (AM9822; Invitrogen) in 1× PBS for 45 min at room temperature, then washed in PBS and left to dry. Once dry, a house made flow cell was attached to the coverslip to allow flow of hybridization reagents. A hybridization mix with 1nM of each oligo in 50% Hybridization Buffer (50% HB: 2 × SSC, 50% Formamide (v/v) (Invitrogen AM9344), 10% Dextran Sulfate (Sigma D8906) in Ultrapure water) was applied to the tissue section. Sections were then incubated for 30 hours at 37°C, washed in 55% Wash Buffer (55% WB: 2 × SSC, 55% Formamide (v/v), 0.1% Triton X-100 (Sigma 93443)) for 30 minutes at 37°C, and then washed twice in 2× SSC at room temperature.

#### Microscope setup and seqFISH imaging

Setup, sequential hybridizations and imaging were performed as previously described(*35*, *74*) with some modifications. Data was captured on a Leica DMi8 microscope equipped with a confocal scanner unit (Yokogawa CSU-W1), fiber-coupled lasers (643, 561, 488 and 405 nm) from CNI, Shanghai Dream Lasers Technology and filter sets from Semrock, a sCMOS camera (Andor Zyla 4.2 Plus), 63 × oil objective lens (Leica 1.40 NA), and a motorized stage (ASI MS2000). A custom-made automated sampler moved designated readout probes in hybridization buffer from a 2.0-ml 96-well plate through a multichannel fluidic valve (IDEX Health & Science EZ1213-820-4) to the custom-made flow cell with a syringe pump (Hamilton Company 63133-01). The syringe pump was used also to move other buffers through the multichannel fluidic valve to the custom-made flow cell. Integration of the imaging and the automated fluidics delivery system was controlled by custom-written scripts in μManager.

For hybridization and imaging, the sample within the custom-made flow cell was first connected to the automated fluidics system on the motorized stage of the microscope. The fields of view (FOVs) were registered using nuclear signals revealed by preincubation in 5 μg ml−1 DAPI (Sigma D8417) in 4× SSC. Imaging was performed with sequential hybridization and imaging routines. The serial hybridization buffer contained three unique readout probes (12.5 nM each) with different fluorophores (Alexa Fluor 647, Cy3B or Alexa Fluor 488) in 10% EC buffer (10% ethylene carbonate (Sigma E26258), 10% dextran sulfate (Sigma D4911) and 4× SSC) and was picked up from a 96-well plate and distributed into the flow cell for a 20 min incubation. After the serial hybridization, the sample was washed with 1 ml of 4× SSCT (4× SSC and 0.1% Triton-X), followed by a washing step with 500 μl of the 12.5% wash buffer. Then, the samples were rinsed with 500 μl of 4× SSC, and stained with 200 μl of the DAPI solution for 60 s to visualize nuclei. Next, anti-bleaching buffer was flown through the sample for imaging. The anti-bleaching buffer was made of 50 mM Tris-HCl pH 8.0 (Invitrogen 15568025), 300 mM NaCl (Invitrogen AM9759), 2× SSC, 3 mM trolox (Sigma 238813), 0.8% D-glucose (Sigma G7528), 1,000-fold diluted catalase (Sigma C3155) and 0.5 mg ml−1 glucose oxidase (Sigma G2133).

Snapshots were acquired with 643-nm, 561-nm, 488-nm and 405-nm fluorescent channels per field of view (FOV). After image acquisition, 1 ml of the 55% wash buffer was flown for 1 min to strip off readout probes, followed by a 5 min incubation before rinsing with 4× SSC. The serial hybridization, imaging and signal extinguishing steps were repeated until all rounds were completed. Blank images displaying only cellular autofluorescence were imaged at the beginning and end of the routines.

Anti-Vcam1 antibody staining was performed as the last step of the hybridizations. Prior to antibody labeling, the tissue was washed multiple times with 1X PBS and a blocking buffer containing 1x PBS (ambion, AM9625) ultra-pure BSA 1% (Ambion, AM 2616) TritonX- 100 0.3% (sigma-Aldrich, 93443) Dextran Sulfate Low MW 0.1% (Sigma-Aldrich, D4911-10G) Salmon Sperm DNA 0.5mg/ml (invitrogen, am9680) was applied and incubated with the sample for 15 minutes at room temperature. Then, primary rabbit anti-Vcam1 antibody (Abcam, ab134047) was introduced at a 1:100 dilution in the blocking buffer and the sample was incubated at room temperature for 1 hr, washed three times in 1X PBS and a secondary anti-rabbit 647 antibody (Invotrogen, A32733) applied at 1:500 dilution for 1 hr at room temperature. Finally, the sample was washed extensively in >2ml of PBS and incubated for 30min with PBS at room temperature to remove excess antibody. Finally, DAPI and anti-bleach buffer were applied before imaging. The same FOVs were then imaged with the antibody label.

#### Immunofluorescence and RNAscope

Immunofluorescence and RNAscope experiments were performed as previously described (*19*). using the following antibodies and probes: mouse IgG2a anti-alpha Smooth Muscle Actin (aSMA, 1:2000, Sigma-Aldrich, A5228), goat anti-Vcam1 (1:200, R&D Systems, AF643), rabbit anti-Vcam1 (1:200, Abcam, ab134047), donkey anti-rabbit AlexaFluor 594 (1:500, Molecular probes, A21207), donkey anti-goat AlexaFlour 594 (1:500, Life Technologies, A11058), chick anti-rat AlexaFluor 647 (1:500, Molecular Probes, A21472). RNAscope Mm-Acta2 (ACD, 319531), RNAscope Mm-Cd4-C3 (ACD, 406841-C3), RNAscope Mm-Cd8a-C2 (ACD, 401681-C2),

RNAscope Probe - Mm-Clcf1-C1 (ACD, 457971), RNAscope Mm-Clec9a-O1 (ACD, 537731), RNAscope Probe - Mm-Cntfr-C3 (ACD, 457981-C3), RNAscope Probe - Mm-Crlf1-C2 (ACD, 446891-C2), RNAscope Probe - Mm-Npr3-C3 (ACD, 502991-C3). Images were acquired on Leica Sp8.

#### Image analysis of seqFISH data

*Image registration:* Image registration was performed using phase cross correlation on DAPI stained images for each FOV. The initial hybridization round was used as reference for estimating required translational shifts.

*Image preprocessing:* Aligned images underwent background subtraction using dilated blank images to further remove unwanted background noise. Post background subtraction, further background removal was performed using a 5 x 5 high pass Gaussian filter, followed by a 3 x 3 low pass Gaussian filter to mitigate hot pixels and make spots more gaussian-like for improved 2D gaussian fitting. Image intensities across channels and serial hybridizations were normalized by 80-99.999% percentile clipping and rescaling between 0-1.

*Spot calls and spot feature generation*: Sub-pixel centroids of identified spots were obtained using DAOStarFinder, a spot picking algorithm from Astropy that performs fast 2D gaussian fits. The full-width half-max (FWHM) was numerically optimized to find the best parameter for spot calling. Features such as flux, peak amplitude, sharpness, bilateral to four-fold symmetry, and symmetry of gaussian fits were recorded and stored for each spot from the DAOStarFinder algorithm. To obtain additional features such as total spot area, a 7×7 bounding box was used to isolate each spot, then a local adaptive threshold using a gaussian kernel was used to obtain the area of the spot.

*Mask generation and spot mapping*: Cell masks were generated using Cellpose 2.0; masks touching the edges of the image were removed to prevent edge bias or undercounting of spots in cells. Additionally, 2 pixels were deleted between two or more masks that touch to prevent spot mixing between cell masks. Spots were mapped to each cell mask to assign a cell IDs.

*Decoding*: Super-resolved, mapped spots were decoded using a support vector machine (SVM) embedded, feature-based symmetrical nearest neighbor algorithm. First, the decoder removed unwanted noise through an SVM model with radial-basis function or polynomial kernel to filter false spots based on their spot characteristics. Post-filtering, each spot was assigned a probability score on the likelihood that a given spot corresponded to a true barcode based on its spot features. Then, the algorithm performed a nearest neighbor radial search for each spot at a distance of 1-2 pixels across barcoding rounds (ignoring spots within the same barcoding round). Searches are performed comparing each round to the other individual rounds in a parallel fashion. When multiple spots fell within the search radius of a given reference spot, each spot was assigned a score based on distance, intensity, and size. The highest scoring spot was chosen and an overall codeword score assigned after picking the best spot for each barcoding round. These scores (including the overall codeword score) are influenced by the number of total neighbors encountered. Each codeword was also assigned an ambiguity score, which is the total number of additional neighbors found over the expected for each barcoding reference round. The probability score (obtained from individual spot probabilities) and the codeword score was used to generate the overall score for each decoded barcode. Each completed barcode also underwent a parity check to distinguish true signal from noise, while also allowing one drop in spot calls. Once the best set of spots was chosen, they were filtered based on the number of times the same set of spots were picked when changing the barcoding round reference, keeping spot sets that appear >= 3 times. If there are any codewords that have overlapping spots, then their overall codeword score was used to pick the best one. If they had the same score, then the codeword with the smallest total distance between spots was used. After the 1st round of decoding, unused spots were resubmitted for additional rounds of decoding (up to 2) to try, and assign left over spots. Decoded spots, including “trues” and “fakes”, were combined to generate a final decoded spots table. These decoded spots were sorted based on their overall codeword score and subsampled to calculate false positive rates (FPR). Generally, the codeword score had a direct relationship with fake spots with low scores corresponding more with fake spots and high scores corresponding to true spots. Once a FPR of 15% was reached, only those subsets of true and fake codes are used for analysis. Once the desired barcodes are obtained, a gene-by-cell matrix is generated for downstream analysis.

*SVM training:* Called spots underwent a quick pass decoding to obtain labels for true and fake spots. The number of fake spots must be 500-500,000 for the SVM classifier to become active. The number of true spots is down-sampled to match the number of fake spots for balanced training. Eighty percent of the data was used for training and 20% for validation. Spot features were normalized using a MinMax Scaler and stored, then the SVM model underwent hyper-parameter tuning of C, gamma, and degree parameters using GridSearchCV with 8-fold cross validation using either a polynomial or radial-basis function kernel. Once the optimal parameters were obtained, the test set underwent the same feature scaling parameters that were used for the training data to gauge out-of-sample performance.

*False Positive Rate:* The false positive rate was calculated as follows:

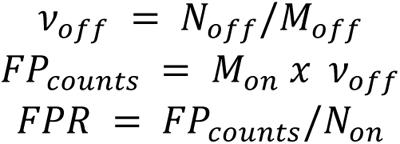

The frequency of off-targets (*ν*_*off*_) was estimated as the number of observed off-target codes (*N*_*off*_) divided by the number of off-target codes in the codebook (*M*_*off*_). Assuming that on-target codes have the same frequency of error as off-target codes, we can multiply the *ν*_*off*_ by the number of on-targets (*M*_*on*_) in the codebook to obtain estimated false positive counts (*FP*_*counts*_). *FP*_*counts*_ were then normalized by the number of observed on-targets (*N*_*on*_) to obtain false positive rate (FPR).

*Code Availability:* Scripts used for pre-processing seqFISH images can be found at https://github.com/CaiGroup/pyfish_tools.

#### Image analysis of serial probes

Images of the serial probes were subjected to the same background subtraction and processing as described above. The RNA spots were then manually thresholder based on visual inspection of the images. The same threshold was used for all experiments

#### Gene expression preprocessing and clustering

The R package Seurat (version 4.1.0) (*75*) was used to process the data. Low quality cells (RNA count ≥ 25 and/or genes/cell ≤ 2) were removed and the data normalized and scaled using the NormalizeData() and ScaleData() function. Principal component analysis was performed using RunPCA(), considering all genes in the dataset. Batch effects were removed using the Harmony algorithm within the Seurat wrapper RunHarmony() (*76*). The functions RunUMAP(), FindNeighbors() and FindClusters() (resolution 0.8) were used for dimensionality reduction and cluster identification, resulting in a total of 31 clusters. The distribution of clusters across samples and imaged positions was examined, and six clusters with a strong position bias caused by technical issues such as imaging in a different focal plane (9452 cells in total) were excluded from downstream analysis, resulting in a final data set of 220,970. Differentially expressed genes between clusters were identified using the FindAllMarkers() function with a Wilcoxon Rank Sum test.

#### ME detection

Detecting the Microenvironments (MEs) was done in the following steps: first, the Euclidean distance between all cells in each sample was calculated. For each cell in the sample, a 30μm radius was defined and the number of cells within this radius which belong to each cell type were counted. Then, the cell-by-neighbor matrix was centered to have a mean of 0, PCA was performed followed by Leiden clustering using Scanpy with sc.pp.neighbors() with n_neighbors = 20 and sc.tl.leiden() with resolution = 0.5. This process resulted in 19 clusters. The results were filtered to remove clusters with < 500 cells total - this resulted in the removal of one cluster. Then the cell type composition in each cluster was evaluated by calculating the mean counts of each cell type within the clusters as in Figure 2D. Two clusters which shared a highly similar cell type composition were manually merged to create a total of 17 clusters (Fig 2D) with 220,753 total cells.

#### NicheNet analysis

NicheNet(*49*) was used to detect signaling between injured cells to the other cell types within the injured niche (ME-5, Figure 3A, Figure S6A). The Seurat implementation NicheNet was used. Only cell types which had ≥ 500 total cells in ME-5 were considered for the analysis. The calculation of top ligand-receptor was run for each of the cell types within ME-5 using nichenet_seuratobj_cluster_de() where the sender was set to be Injured EC, the receiver_reference = cell type in Control and receiver_effected = cell type in AKI. The top 10 ligand - receptor pairs were chosen as pairs with the highest ‘weight’ term out of all pairs that NicheNet identified. Figure S6A plots the mean expression of the top ligand-receptor pairs for all cell types in ME-5.

#### Fibroblasts differential expression analysis

To reduce noise, we first calculated the average expression of all genes across all cell types and z-scored average expression values. All genes with a z-score < 0.1 within fibroblasts were removed from differential expression analysis. We then identified the differentially expressed genes between fibroblasts within the different MEs using the FindAllMarkers() function in Seurat with the following parameters: min.pct = 0.1,logfc.threshold = 0.25,only.pos = T. Figure 4B shows the average expression of the genes detected in each of the AKI enriched or AKI depleted MEs - as calculated by the enrichment of each ME in the control or AKI using a one sided students’ t-test (significance values shown in Figure 2B). To calculate the score of each gene set on sn-RNAseq data presented in Figure 4D the function AddModuleScore() was used with the differentially expressed genes as features, to apply a score to fibroblasts in the sn-RNAseq data as identified by the authors (*15*). Score values were averaged for each mouse in the data.

#### T cell reference mapping

For T cell reference mapping A reference atlas of T cell subsets was used (*68*) reference RDS object can be found here: https://figshare.com/articles/dataset/ProjecTILs_murine_reference_atlas_of_tumor-infiltrating_T_cells_version_1/12478571/2. RunUMAP() was run on the reference object to return the UMAP model and then reference mapping was done using 1121 genes present in both datasets. Anchors were found with the FindTransferAnchors() functions with normalization.method = “LogNormalize” and reference.reduction = “pca”. Then, MapQuery() was done with the detected anchors with reference.reduction = “pca” and reduction.model = “umap”. Reference cell types which were mapped with ≤ 100 cells in total were removed.

## Supplementary Figures

**Figure S1:**
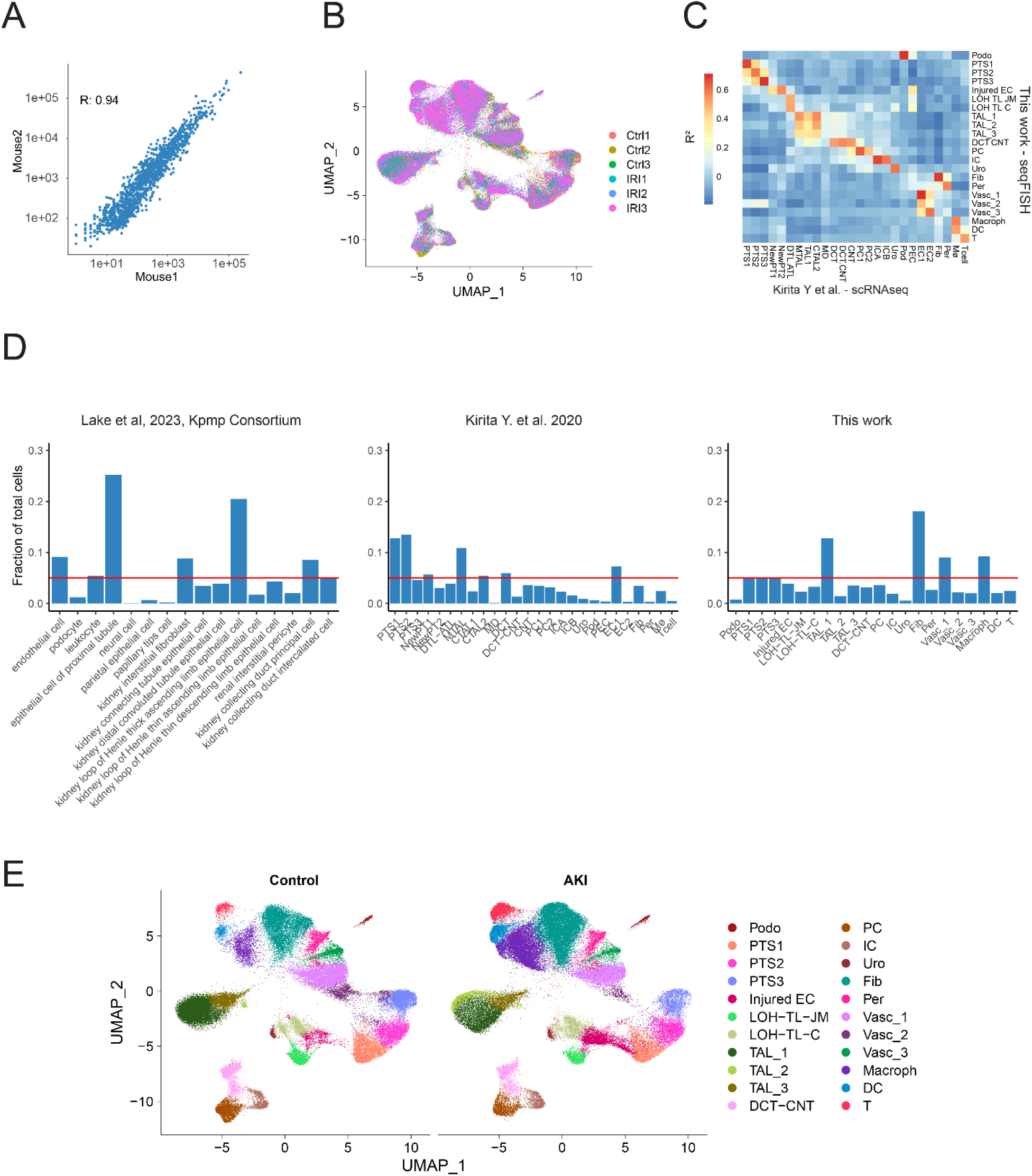
Validation of seqFISH results. A) Pseudo Bulk seqFISH gene expression of two control mice. B) Umap representation of the data color coded by sample. C) Correlation to snRNAseq data. Rsqr showing the Pearson correlation of the average gene expression for all genes common to both datasets in each cell type. D) Fraction of each cell type out of the total cells detected in three datasets: left - human scRNAseq data taken from Lake et al., middle - mouse scRNAseq from Kirita et al. and right - this work. Horizontal line labels 5% datapoint in each plot. E) Umap representation of the single cell data separating cells from control and AKI mice.

**Figure S2:**
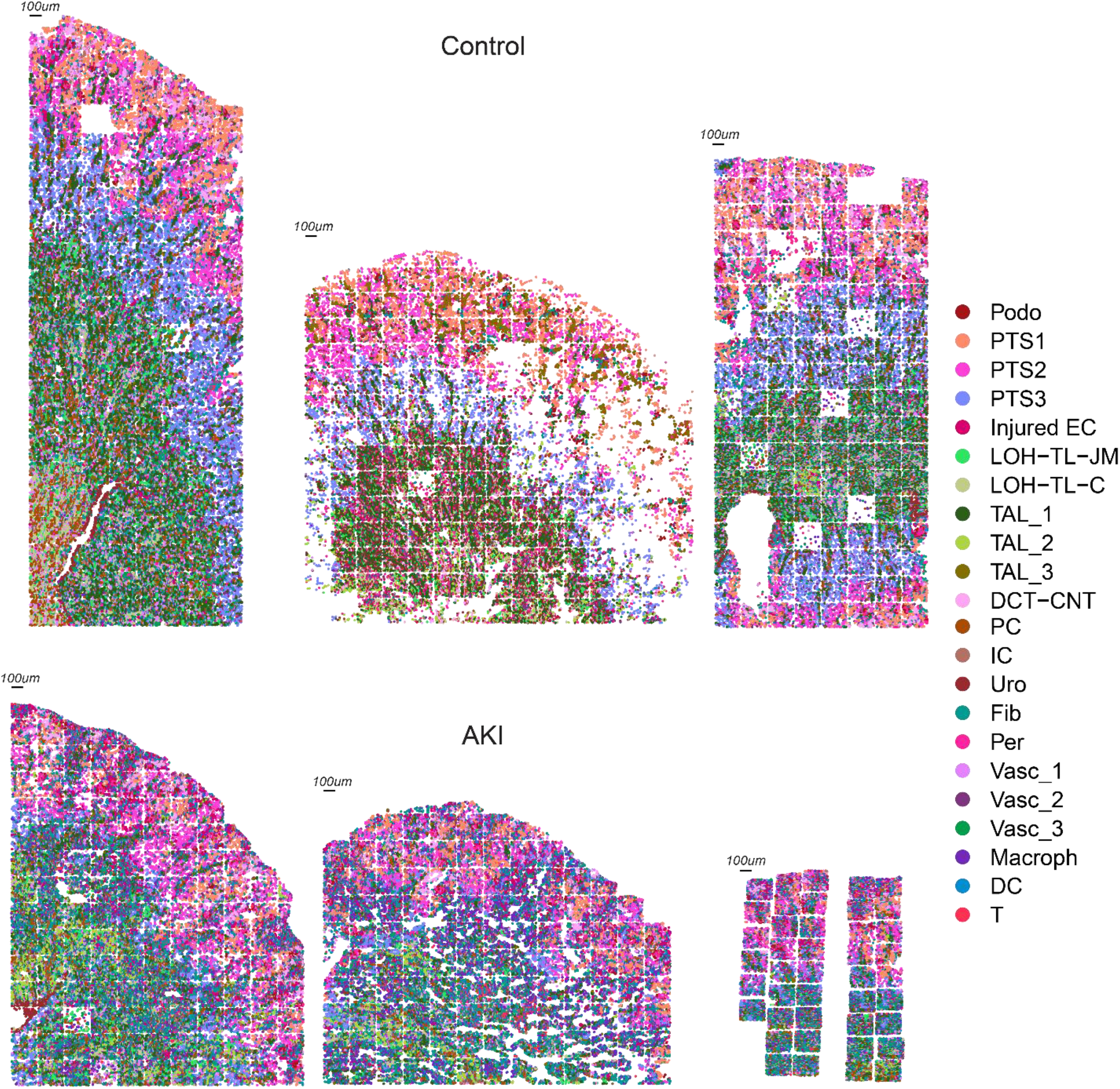
cell type composition of all samples.

**Figure S3:**
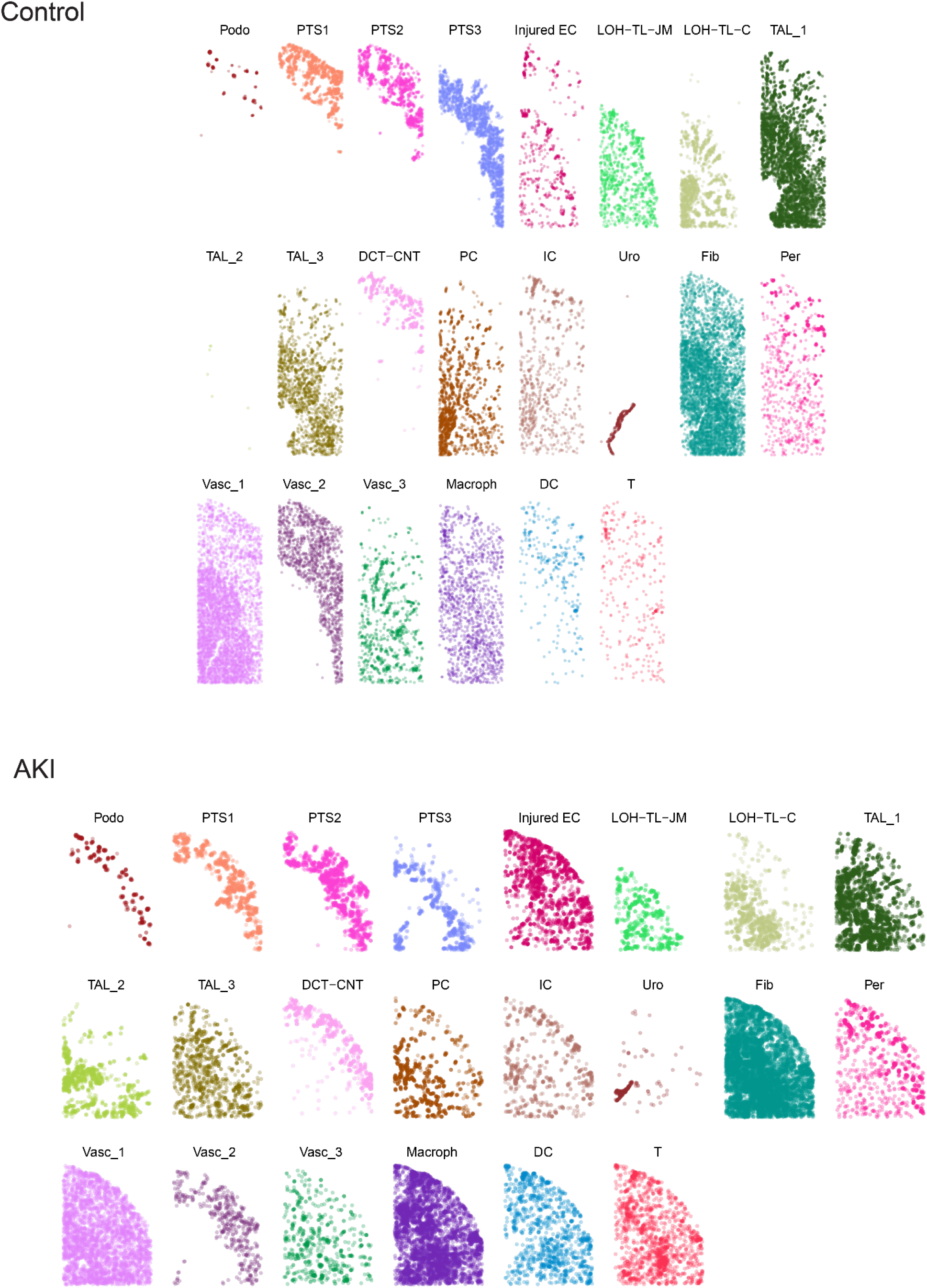
Cell type composition of one control and one AKI sample.

**Figure S4:**
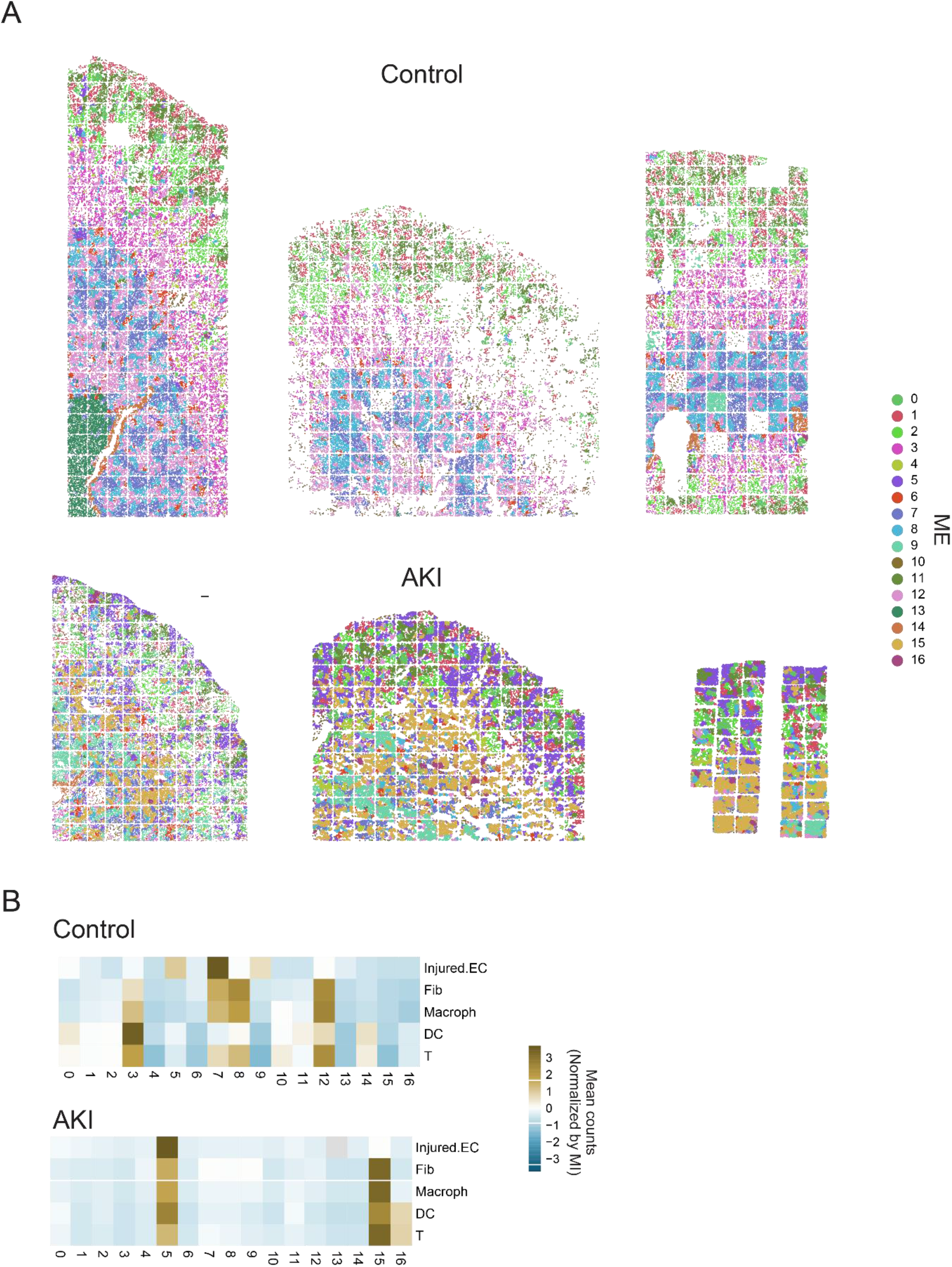
ME composition of control and AKI samples: A) ME composition of all six control and AKI samples. B) Cell type distribution of AKI-enriched cells within the different MEs within control and AKI samples.

**Figure S5:**
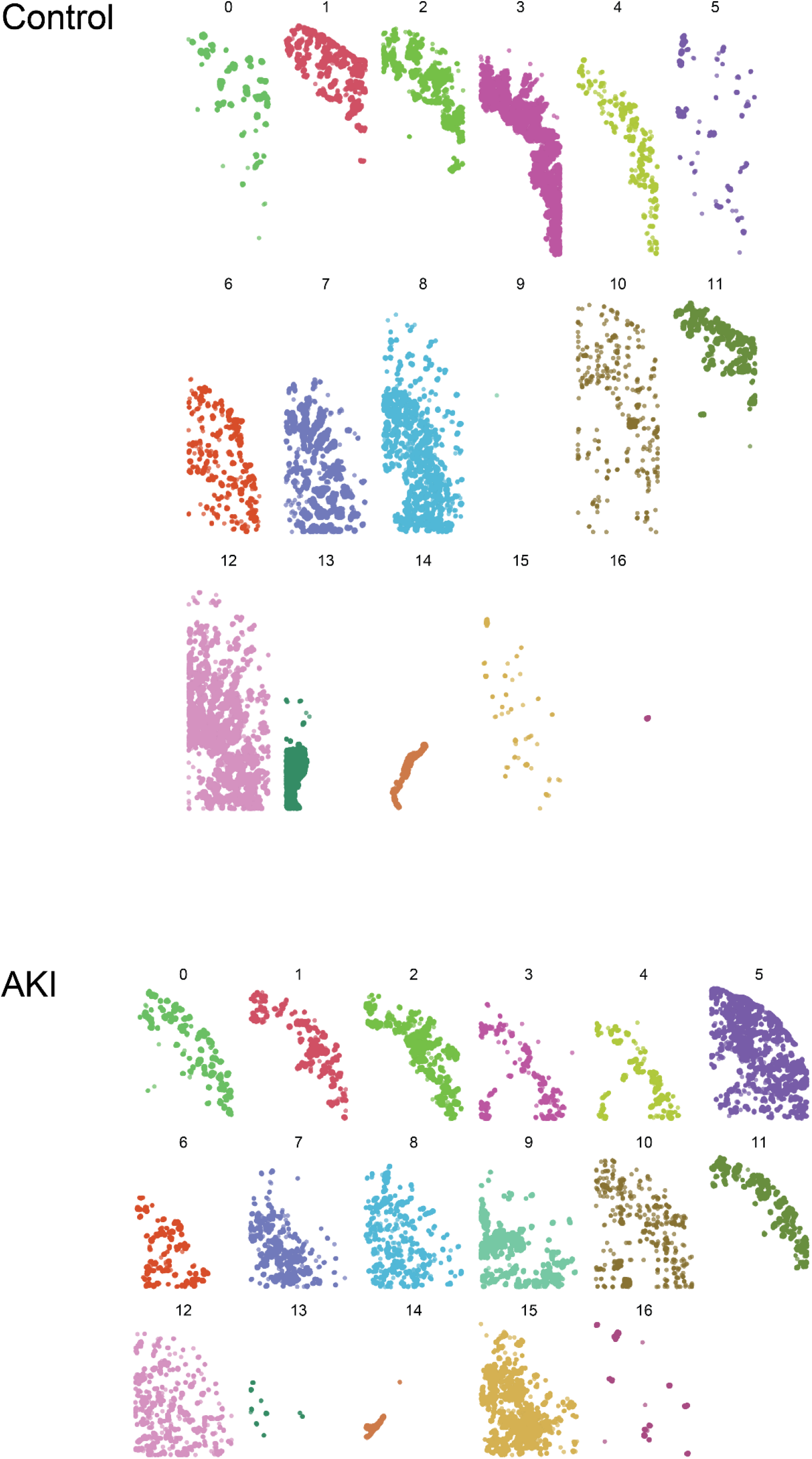
ME composition in one control and one AKI samples

**Figure S6:**
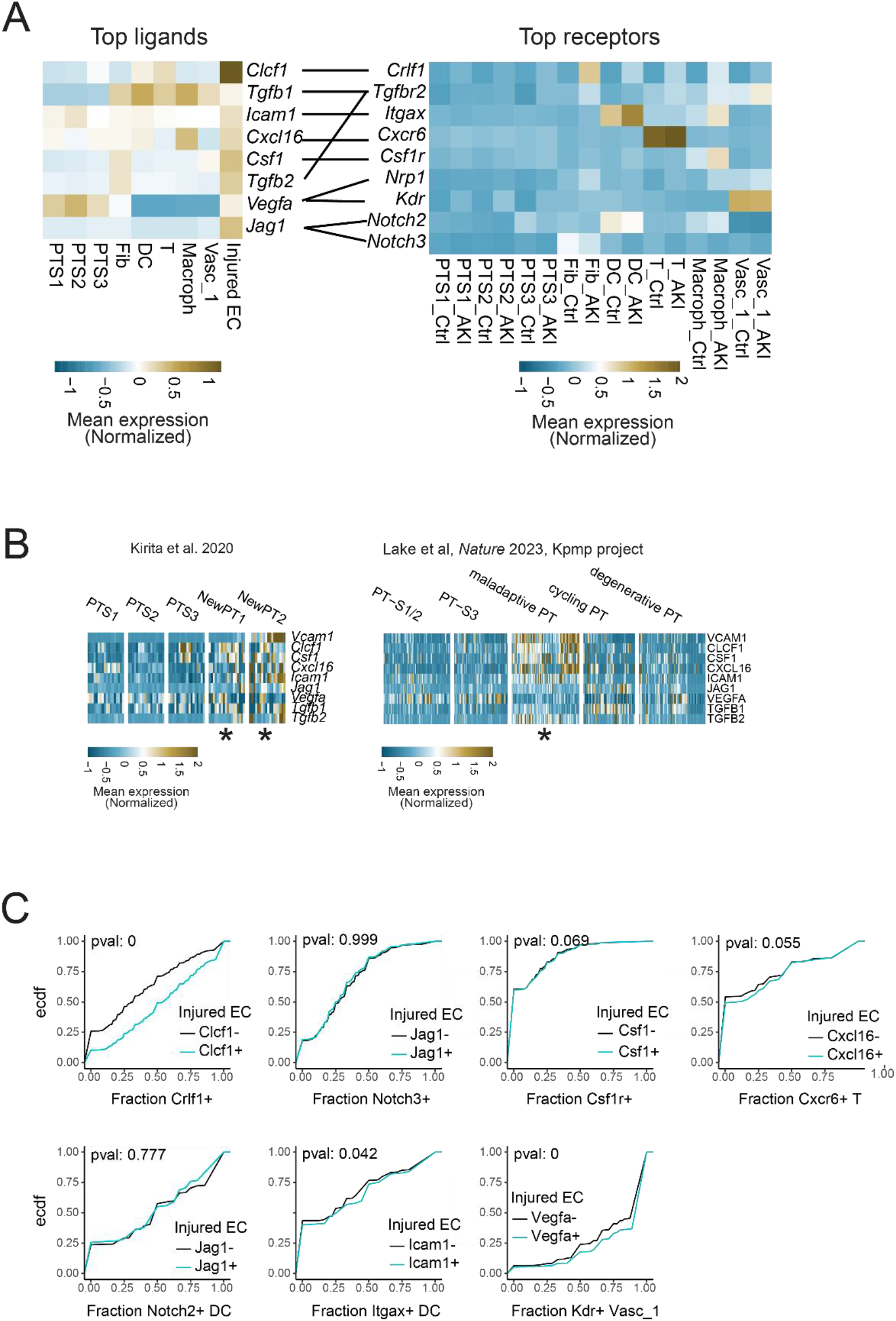
Ligand - Receptor expression within ME-5: A) Average expression of top ligands in cell types found within ME-5 (left) and expression of the top receptors within the same cell types, in either control or AKI samples (right). Expression is averaged for each cell type within and outside of ME-5. Lines indicate ligand and receptor pairs. B) Left – Expression of the top ligands as in A, in PT cells detected within a snRNAseq dataset from Kirita et al. Gene expression was normalized per sample, heatmap is showing the mean expression of the indicated cell types for all samples. [NewPT1, NewPT2: populations of injured epithelial cells appearing post AKI, noted with *]. Right - Expression of the same ligands in PT cells within the human Kpmp scRNAseq dataset. Gene expression was normalized per sample, and heatmap is showing the mean expression of the indicated cell types for all samples. [PT-S1/2, PT-S3: Proximal tubule cells segments 1/2 and 3, respectively. Maladaptive PT: Proximal tubule cells appearing following AKI. Cycling PT: Proximal tubule cells undergoing division, degenerative PT: PT cells unable to execute repair. * denotes injured epithelial cells]. C) ECDF as in main figure 3. For all ligand-receptor pairs within the cell types that were found in ME-5

**Figure S7.**
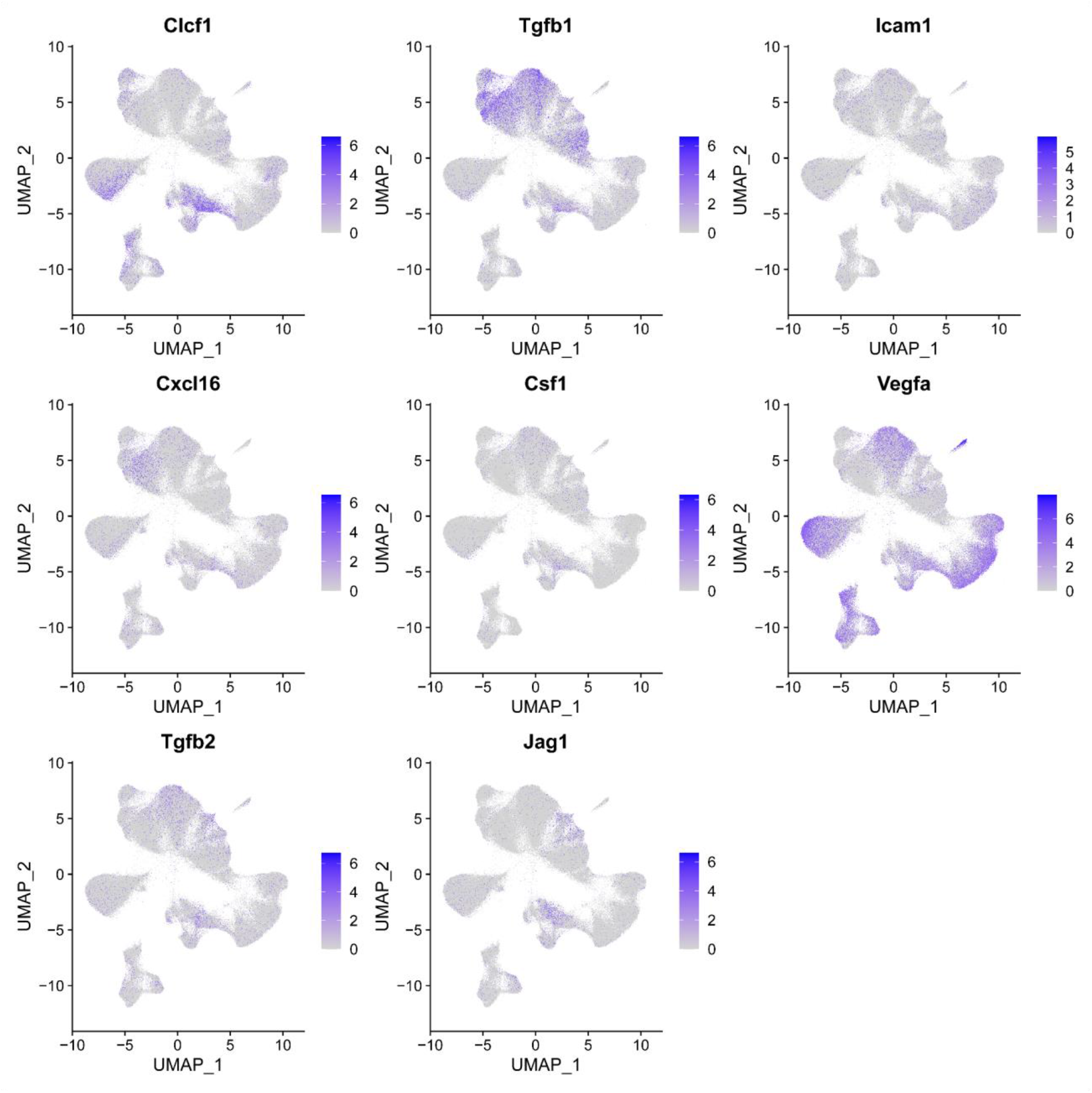
top ligands on the cell type Umap.

**Figure S8:**
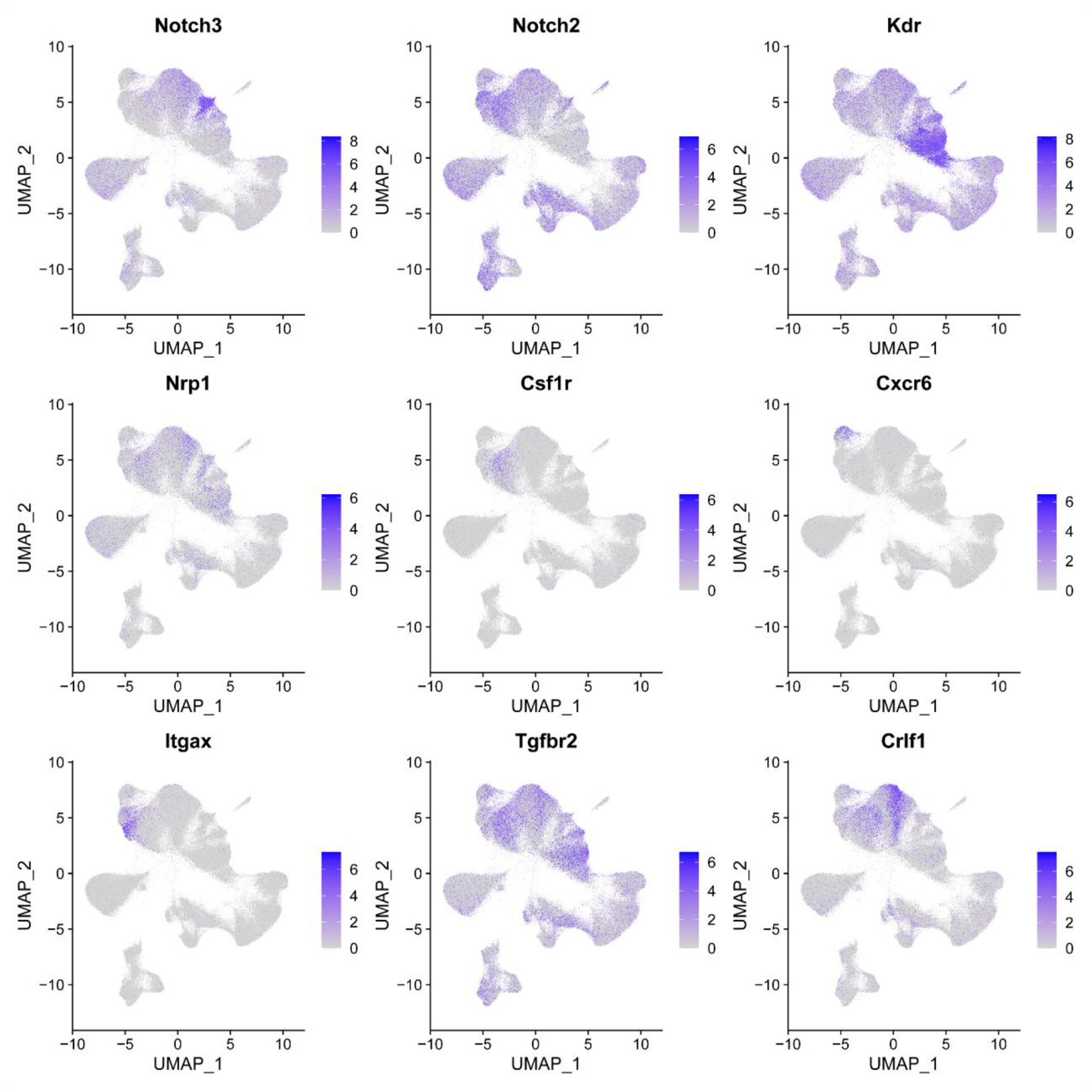
Top receptors on the cell type Umap.

**Figure S9:**
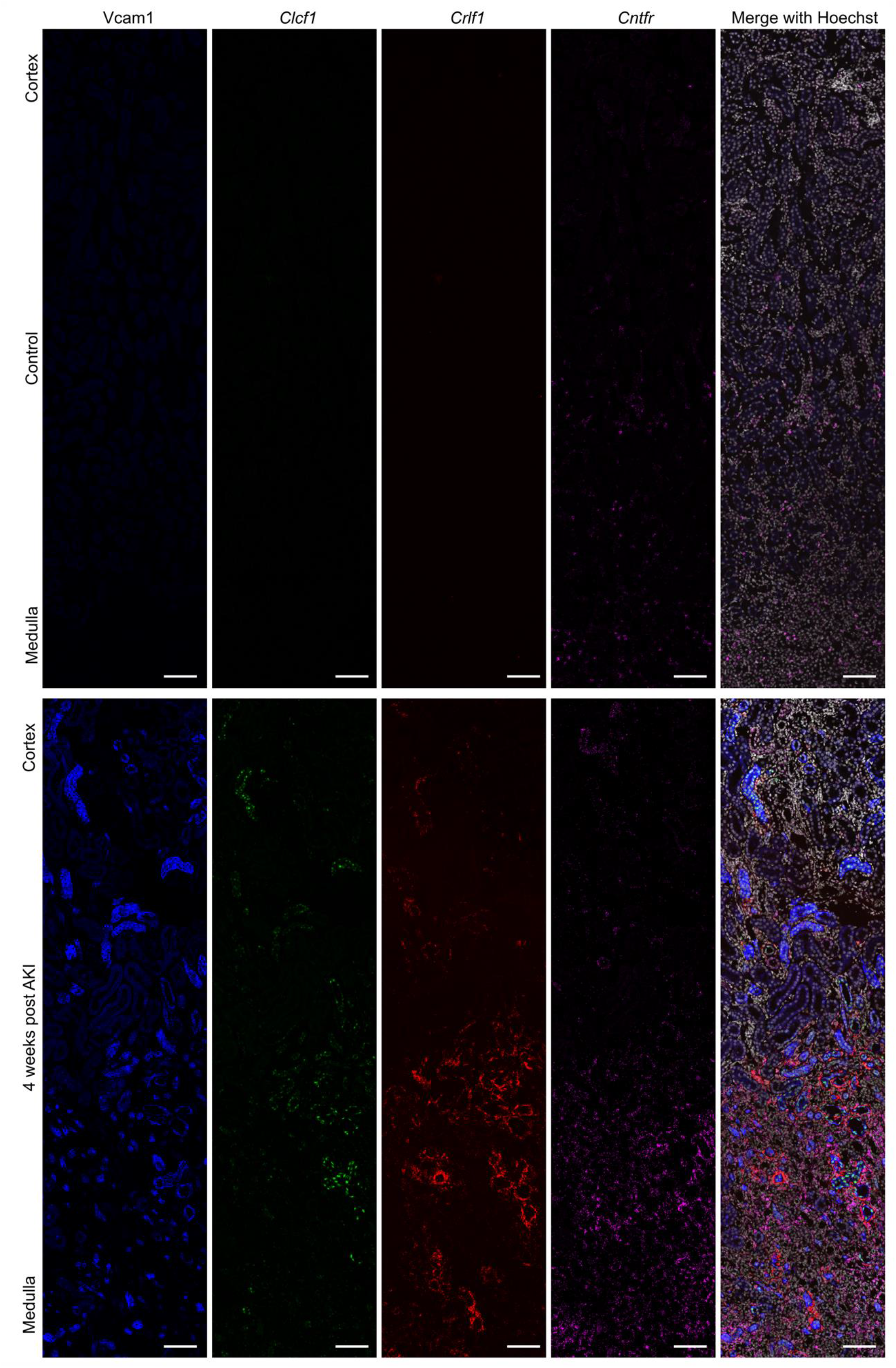
Validation of expression pattern and spatial localization of *Clcf1* and *Crlf1*. (A) Validation of observed expression changes and spatial colocalization of *Clcf1* and *Crlf1* after injury by a combination of immunofluorescence staining (Vcam1) and RNAscope *in situ* hybridization (*Clcf1, Crlf1*, *Cntfr*). Scale bar = 100 µm.

**Figure S10:**
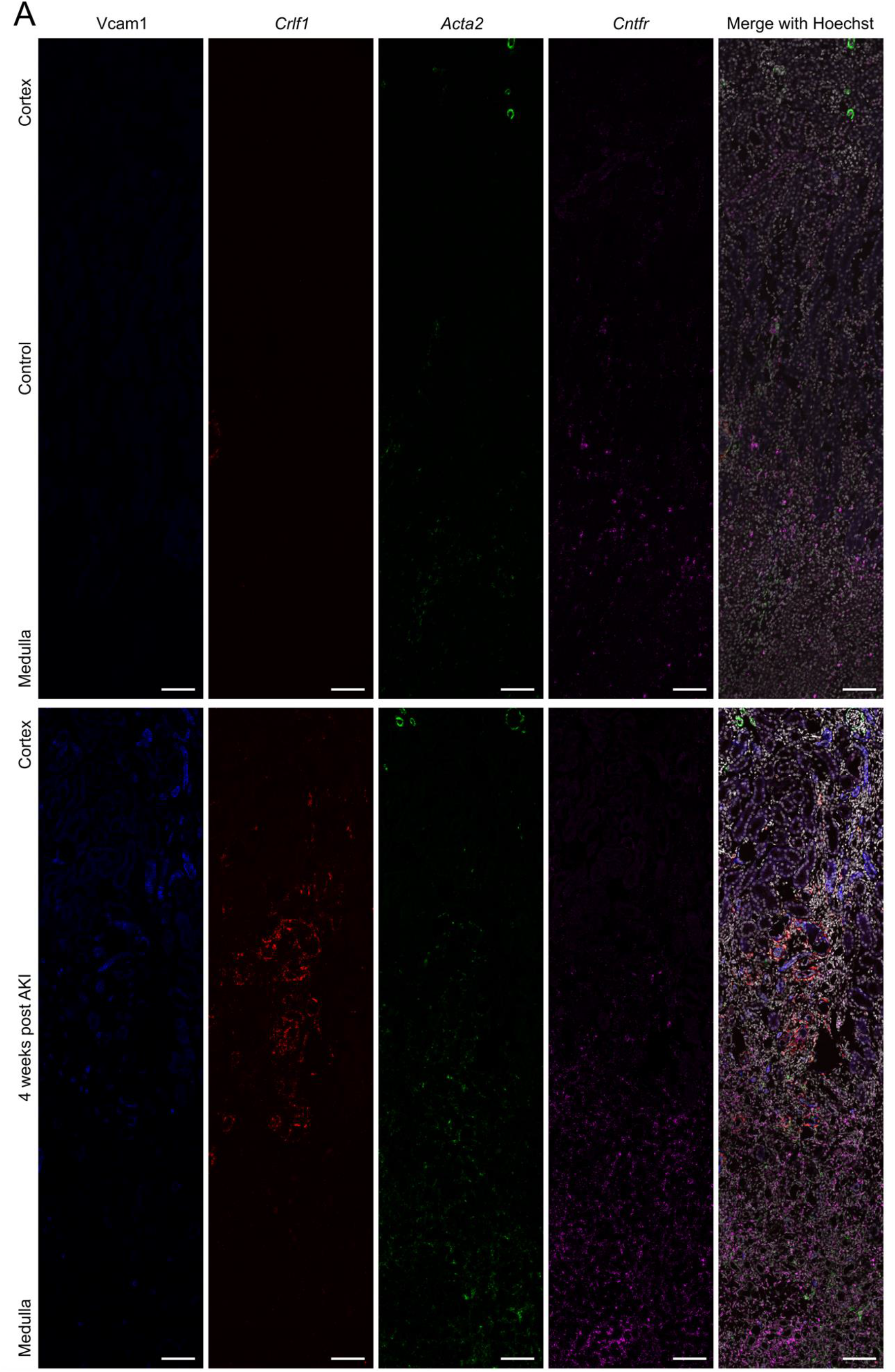
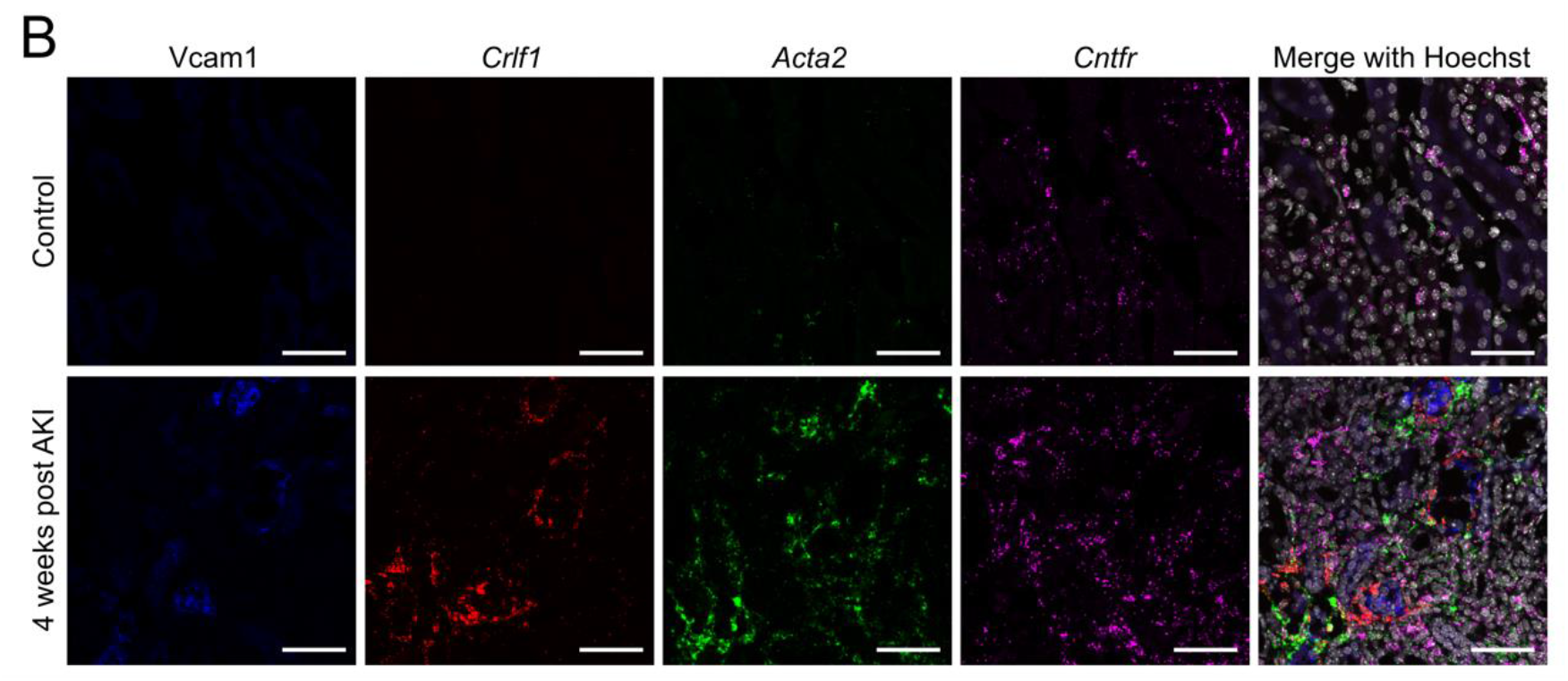
C*rlf1* marks fibroblasts around Vcam1+ injured epithelial cells. A combination of immunofluorescence staining (Vcam1) and RNAscope *in situ* hybridization (*Crlf1*, *Acta2*, *Cntfr*) was used to validate *Crlf1* as a marker of fibroblasts spatially associated with Vcam1+ injured epithelial cells. Scale bar = 100 µm. (B) Zoom in of (A). Scale bar = 50µm.

**Figure S11:**
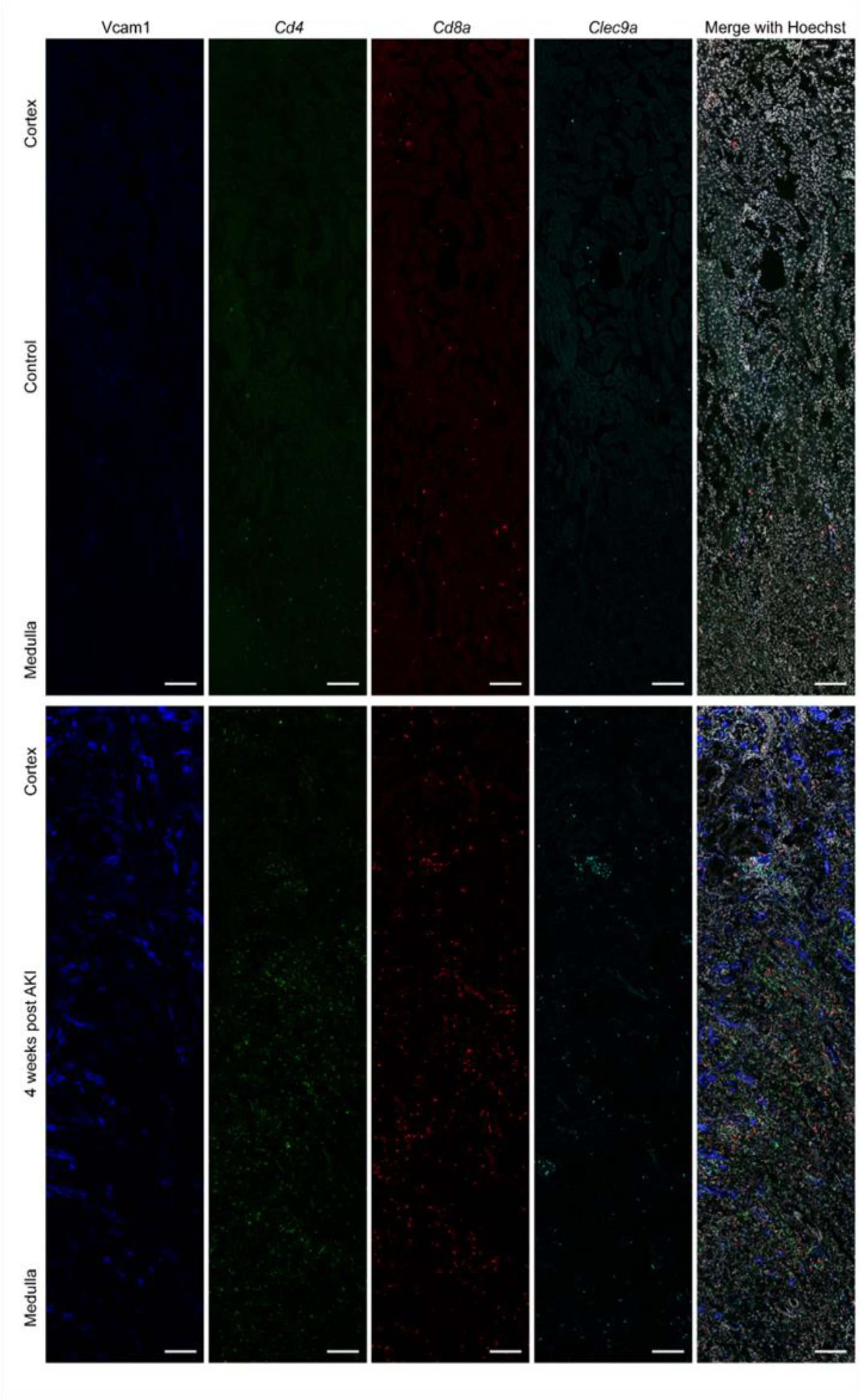
Expression of *Cd4*, *Cd8a* and *Clec9* on control and AKI kidneys. A combination of immunofluorescence staining (Vcam1) and RNAscope *in situ* hybridization (*Cd4, Cd8a, Clec9*) was used to assess the spatial distribution of CD4 and CD8 T cells as well as dendritic cells. Scale bar = 100 µm.

**Figure S12:**
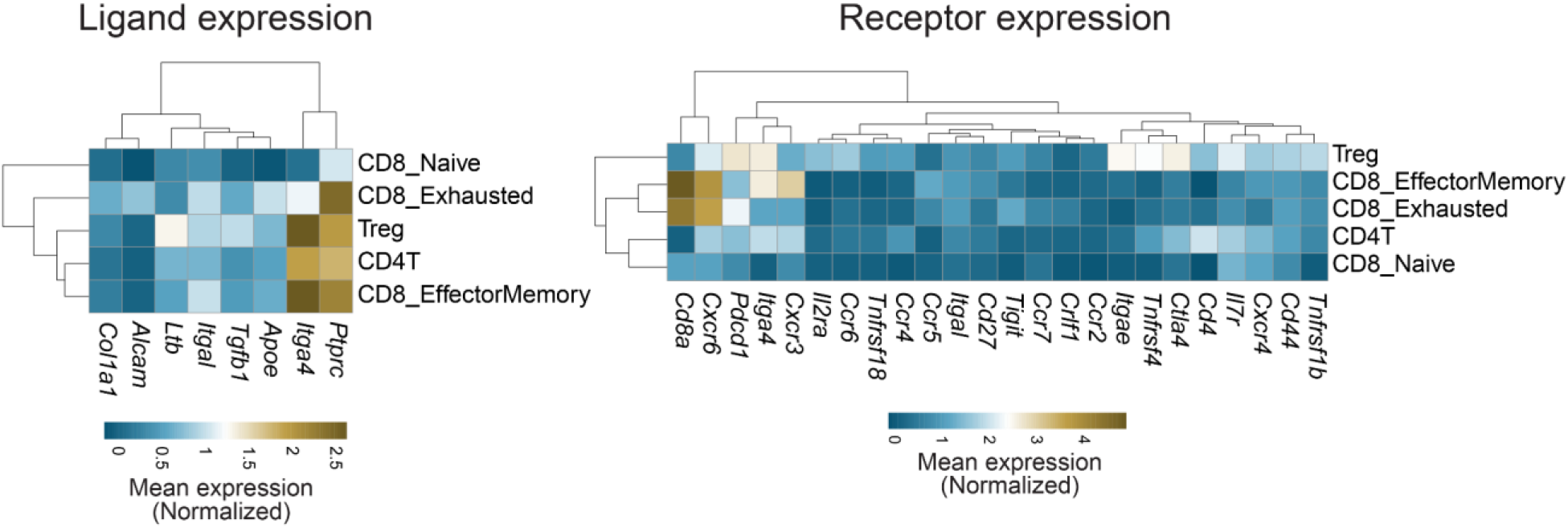
Ligand receptor expression in T cell subsets.

## References

1. W. Su, R. Cao, X.-Y. Zhang, Y. Guan, Aquaporins in the kidney: physiology and pathophysiology. Am. J. Physiol. Renal Physiol. 318, F193–F203 (2020).

2. A. S. L. Yu, G. M. Chertow, V. A. Luyckx, P. A. Marsden, K. Skorecki, M.W. Taal, Brenner & Rector’s the Kidney (2020).

3. A. Ransick, N. O. Lindström, J. Liu, Q. Zhu, J.-J. Guo, G. F. Alvarado, A. D. Kim, H. G. Black, J. Kim, A. P. McMahon, Single-Cell Profiling Reveals Sex, Lineage, and Regional Diversity in the Mouse Kidney. Dev. Cell. 51, 399–413.e7 (2019).

4. Acute kidney injury. Lancet. 394, 1949–1964 (2019).

5 International Society of Nephrology’s 0by25 initiative for acute kidney injury (zero preventable deaths by 2025): a human rights case for nephrology. Lancet. 385, 2616–2643 (2015).

6. J. A. Kellum, P. Romagnani, G. Ashuntantang, C. Ronco, A. Zarbock, H.-J. Anders, Acute kidney injury. Nature Reviews Disease Primers. 7, 1–17 (2021).

7. P. Romagnani, G. Remuzzi, R. Glassock, A. Levin, K. J. Jager, M. Tonelli, Z. Massy, C. Wanner, H.-J. Anders, Chronic kidney disease. Nature Reviews Disease Primers. 3, 1–24 (2017).

8 C. Kuppe, M. M. Ibrahim, J. Kranz, X. Zhang, S. Ziegler, J. Perales-Patón, J. Jansen, K. C. Reimer, J. R. Smith, R. Dobie, J. R. Wilson-Kanamori, M. Halder, Y. Xu, N. Kabgani, N. Kaesler, M. Klaus, L. Gernhold, V. G. Puelles, T. B. Huber, P. Boor, S. Menzel, R. M. Hoogenboezem, E. M. J. Bindels, J. Steffens, J. Floege, R. K. Schneider, J. Saez-Rodriguez, N. C. Henderson, R. Kramann, Decoding myofibroblast origins in human kidney fibrosis. Nature. 589, 281–286 (2020).

9 Forecasting life expectancy, years of life lost, and all-cause and cause-specific mortality for 250 causes of death: reference and alternative scenarios for 2016–40 for 195 countries and territories. Lancet. 392, 2052–2090 (2018).

10. L. S. Chawla, P. L. Kimmel, Acute kidney injury and chronic kidney disease: an integrated clinical syndrome. Kidney Int. 82, 516–524 (2012).

11. R. Witzgall, D. Brown, C. Schwarz, J. V. Bonventre, Localization of proliferating cell nuclear antigen, vimentin, c-Fos, and clusterin in the postischemic kidney. Evidence for a heterogenous genetic response among nephron segments, and a large pool of mitotically active and dedifferentiated cells. J. Clin. Invest. 93, 2175–2188 (1994).

12. T. Kusaba, M. Lalli, R. Kramann, A. Kobayashi, B. D. Humphreys, Differentiated kidney epithelial cells repair injured proximal tubule. Proc. Natl. Acad. Sci. U. S. A. 111, 1527–1532 (2014).

13. L. M. S. Gerhardt, J. Liu, K. Koppitch, P. E. Cippà, A. P. McMahon, Single-nuclear transcriptomics reveals diversity of proximal tubule cell states in a dynamic response to acute kidney injury. Proc. Natl. Acad. Sci. U. S. A. 118 (2021), doi:10.1073/pnas.2026684118.

14. S. Ide, Y. Kobayashi, K. Ide, S. A. Strausser, K. Abe, S. Herbek, L. L. O’Brien, S. D. Crowley, L. Barisoni, A. Tata, P. R. Tata, T. Souma, Ferroptotic stress promotes the accumulation of pro-inflammatory proximal tubular cells in maladaptive renal repair. Elife. 10 (2021), doi:10.7554/eLife.68603.

15. Y. Kirita, H. Wu, K. Uchimura, P. C. Wilson, B. D. Humphreys, Cell profiling of mouse acute kidney injury reveals conserved cellular responses to injury. Proc. Natl. Acad. Sci. U. S. A. 117, 15874–15883 (2020).

16. R. Kramann, M. Tanaka, B. D. Humphreys, Fluorescence microangiography for quantitative assessment of peritubular capillary changes after AKI in mice. J. Am. Soc. Nephrol. 25, 1924–1931 (2014).

17. E. E. Dixon, H. Wu, Y. Muto, P. C. Wilson, B. D. Humphreys, Spatially Resolved Transcriptomic Analysis of Acute Kidney Injury in a Female Murine Model. J. Am. Soc. Nephrol. 33, 279–289 (2022).

18. E. Ó hAinmhire, B. D. Humphreys, Fibrotic Changes Mediating Acute Kidney Injury to Chronic Kidney Disease Transition. Nephron. 137, 264–267 (2017).

19 L. M. S. Gerhardt, K. Koppitch, J. van Gestel, J. Guo, S. Cho, H. Wu, Y. Kirita, B. D. Humphreys, A. P. McMahon, Lineage Tracing and Single-Nucleus Multiomics Reveal Novel Features of Adaptive and Maladaptive Repair after Acute Kidney Injury. J. Am. Soc. Nephrol. 34, 554–571 (2023).

20. Y. Sato, P. Boor, S. Fukuma, B. M. Klinkhammer, H. Haga, O. Ogawa, J. Floege, M. Yanagita, Developmental stages of tertiary lymphoid tissue reflect local injury and inflammation in mouse and human kidneys. Kidney Int. 98, 448–463 (2020).

21. B. R. Conway, E. D. O’Sullivan, C. Cairns, J. O’Sullivan, D. J. Simpson, A. Salzano, K. Connor, P. Ding, D. Humphries, K. Stewart, O. Teenan, R. Pius, N. C. Henderson, C. Bénézech, P. Ramachandran, D. Ferenbach, J. Hughes, T. Chandra, L. Denby, Kidney Single-Cell Atlas Reveals Myeloid Heterogeneity in Progression and Regression of Kidney Disease. J. Am. Soc. Nephrol. 31, 2833–2854 (2020).

22. Y. Sato, K. Silina, M. van den Broek, K. Hirahara, M. Yanagita, The roles of tertiary lymphoid structures in chronic diseases. Nat. Rev. Nephrol., 1–13 (2023).

23. R. Huang, P. Fu, L. Ma, Kidney fibrosis: from mechanisms to therapeutic medicines. Signal Transduct Target Ther. 8, 129 (2023).

24. L. Li, H. Fu, Y. Liu, The fibrogenic niche in kidney fibrosis: components and mechanisms. Nat. Rev. Nephrol. 18, 545–557 (2022).

25. C. Hinze, C. Kocks, J. Leiz, N. Karaiskos, A. Boltengagen, S. Cao, C. M. Skopnik, J. Klocke, J.-H. Hardenberg, H. Stockmann, I. Gotthardt, B. Obermayer, L. Haghverdi, E. Wyler, M. Landthaler, S. Bachmann, A. C. Hocke, V. Corman, J. Busch, W. Schneider, N. Himmerkus, M. Bleich, K.-U. Eckardt, P. Enghard, N. Rajewsky, K. M. Schmidt-Ott, Single-cell transcriptomics reveals common epithelial response patterns in human acute kidney injury. Genome Med. 14, 103 (2022).

26. M. S. Balzer, T. Doke, Y.-W. Yang, D. L. Aldridge, H. Hu, H. Mai, D. Mukhi, Z. Ma, R. Shrestha, M. B. Palmer, C. A. Hunter, K. Susztak, Single-cell analysis highlights differences in druggable pathways underlying adaptive or fibrotic kidney regeneration. Nat. Commun. 13, 4018 (2022).

27. J. Liu, S. Kumar, E. Dolzhenko, G. F. Alvarado, J. Guo, C. Lu, Y. Chen, M. Li, M. C. Dessing, R. K. Parvez, P. E. Cippà, A. M. Krautzberger, G. Saribekyan, A. D. Smith, A. P. McMahon, Molecular characterization of the transition from acute to chronic kidney injury following ischemia/reperfusion. JCI Insight. 2 (2017), doi:10.1172/jci.insight.94716.

28. B. B. Lake, R. Menon, S. Winfree, Q. Hu, R. M. Ferreira, K. Kalhor, D. Barwinska, E. A. Otto, M. Ferkowicz, D. Diep, N. Plongthongkum, A. Knoten, S. Urata, A. S. Naik, S. Eddy, B. Zhang, Y. Wu, D. Salamon, J. C. Williams, X. Wang, K. S. Balderrama, P. Hoover, E. Murray, A. Vijayan, F. Chen, S. S. Waikar, S. Rosas, F. P. Wilson, P. M. Palevsky, K. Kiryluk, J. R. Sedor, R. D. Toto, C. Parikh, E. H. Kim, E. Z. Macosko, P. V. Kharchenko, J. P. Gaut, J. B. Hodgin, M. T. Eadon, P. C. Dagher, T. M. El-Achkar, K. Zhang, M. Kretzler, S. Jain, for the KPMP consortium, An atlas of healthy and injured cell states and niches in the human kidney. Nature (2023), doi:10.1038/s41586-023-05769-3.

29. A. Zuk, J. V. Bonventre, Acute Kidney Injury. Annu. Rev. Med. 67, 293–307 (2016).

30. D. A. Ferenbach, J. V. Bonventre, Mechanisms of maladaptive repair after AKI leading to accelerated kidney ageing and CKD. Nat. Rev. Nephrol. 11, 264–276 (2015).

31. G. Canaud, J. V. Bonventre, Cell cycle arrest and the evolution of chronic kidney disease from acute kidney injury. Nephrol. Dial. Transplant. 30, 575–583 (2015).

32. J. Hansen, R. Sealfon, R. Menon, M. T. Eadon, B. B. Lake, B. Steck, K. Anjani, S. Parikh, T. K. Sigdel, G. Zhang, D. Velickovic, D. Barwinska, T. Alexandrov, D. Dobi, P. Rashmi, E. A. Otto, M. Rivera, M. P. Rose, C. R. Anderton, J. P. Shapiro, A. Pamreddy, S. Winfree, Y. Xiong, Y. He, I. H. de Boer, J. B. Hodgin, L. Barisoni, A. S. Naik, K. Sharma, M. M. Sarwal, K. Zhang, J. Himmelfarb, B. Rovin, T. M. El-Achkar, Z. Laszik, J. C. He, P. C. Dagher, M. T. Valerius, S. Jain, L. M. Satlin, O. G. Troyanskaya, M. Kretzler, R. Iyengar, E. U. Azeloglu, Kidney Precision Medicine Project, A reference tissue atlas for the human kidney. Sci Adv. 8, eabn4965 (2022).

33. F. Schreibing, R. Kramann, Mapping the human kidney using single-cell genomics. Nat. Rev. Nephrol. 18, 347–360 (2022).

34. S. Shah, E. Lubeck, W. Zhou, L. Cai, seqFISH Accurately Detects Transcripts in Single Cells and Reveals Robust Spatial Organization in the Hippocampus. Neuron. 94, 752–758.e1 (2017).

35. C.-H. L. Eng, M. Lawson, Q. Zhu, R. Dries, N. Koulena, Y. Takei, J. Yun, C. Cronin, C. Karp, G.-C. Yuan, L. Cai, Transcriptome-scale super-resolved imaging in tissues by RNA seqFISH. Nature. 568, 235–239 (2019).

36. K. H. Chen, A. N. Boettiger, J. R. Moffitt, S. Wang, X. Zhuang, RNA imaging. Spatially resolved, highly multiplexed RNA profiling in single cells. Science. 348, aaa6090 (2015).

37. X. Wang, W. E. Allen, M. A. Wright, E. L. Sylwestrak, N. Samusik, S. Vesuna, K. Evans, C. Liu, C. Ramakrishnan, J. Liu, G. P. Nolan, F.-A. Bava, K. Deisseroth, Three-dimensional intact-tissue sequencing of single-cell transcriptional states. Science. 361 (2018), doi:10.1126/science.aat5691.

38. S. Alon, D. R. Goodwin, A. Sinha, A. T. Wassie, F. Chen, E. R. Daugharthy, Y. Bando, A. Kajita, A. G. Xue, K. Marrett, R. Prior, Y. Cui, A. C. Payne, C.-C. Yao, H.-J. Suk, R. Wang, C.-C. J. Yu, P. Tillberg, P. Reginato, N. Pak, S. Liu, S. Punthambaker, E. P. R. Iyer, R. E. Kohman, J. A. Miller, E. S. Lein, A. Lako, N. Cullen, S. Rodig, K. Helvie, D. L. Abravanel, N. Wagle, B. E. Johnson, J. Klughammer, M. Slyper, J. Waldman, J. Jané-Valbuena, O. Rozenblatt-Rosen, A. Regev, IMAXT Consortium, G. M. Church, A. H. Marblestone, E. S. Boyden, Expansion sequencing: Spatially precise in situ transcriptomics in intact biological systems. Science. 371 (2021), doi:10.1126/science.aax2656.

39. S. Shah, E. Lubeck, W. Zhou, L. Cai, In Situ Transcription Profiling of Single Cells Reveals Spatial Organization of Cells in the Mouse Hippocampus. Neuron. 92, 342–357 (2016).

40. T. Lohoff, S. Ghazanfar, A. Missarova, N. Koulena, N. Pierson, J. A. Griffiths, E. S. Bardot, C.-H. L. Eng, R. C. V. Tyser, R. Argelaguet, C. Guibentif, S. Srinivas, J. Briscoe, B. D. Simons, A.-K. Hadjantonakis, B. Göttgens, W. Reik, J. Nichols, L. Cai, J. C. Marioni, Integration of spatial and single-cell transcriptomic data elucidates mouse organogenesis. Nat. Biotechnol. 40, 74–85 (2022).

41. Q. Zhu, S. Shah, R. Dries, L. Cai, G.-C. Yuan, Identification of spatially associated subpopulations by combining scRNAseq and sequential fluorescence in situ hybridization data. Nat. Biotechnol. (2018), doi:10.1038/nbt.4260.

42. S. Gaedcke, J. Sinning, O. Dittrich-Breiholz, H. Haller, I. Soerensen-Zender, C. M. Liao, A. Nordlohne, P. Sen, S. von Vietinghoff, D. S. DeLuca, R. Schmitt, Single cell versus single nucleus: transcriptome differences in the murine kidney after ischemia-reperfusion injury. Am. J. Physiol. Renal Physiol. 323, F171–F181 (2022).

43. E. E. Dixon, H. Wu, E. Sulvarán-Guel, J. Guo, B. D. Humphreys, Spatially resolved transcriptomics and the kidney: many opportunities. Kidney Int. 102, 482–491 (2022).

44. E. Denisenko, B. B. Guo, M. Jones, R. Hou, L. de Kock, T. Lassmann, D. Poppe, O. Clément, R. K. Simmons, R. Lister, A. R. R. Forrest, Systematic assessment of tissue dissociation and storage biases in single-cell and single-nucleus RNA-seq workflows. Genome Biol. 21, 130 (2020).

45. H. Wu, Y. Kirita, E. L. Donnelly, B. D. Humphreys, Advantages of Single-Nucleus over Single-Cell RNA Sequencing of Adult Kidney: Rare Cell Types and Novel Cell States Revealed in Fibrosis. J. Am. Soc. Nephrol. 30, 23–32 (2019).

46. H. Scholz, F. J. Boivin, K. M. Schmidt-Ott, S. Bachmann, K.-U. Eckardt, U. I. Scholl, P. B. Persson, Kidney physiology and susceptibility to acute kidney injury: implications for renoprotection. Nat. Rev. Nephrol. 17, 335–349 (2021).

47. L. Chen, C.-L. Chou, M. A. Knepper, Targeted Single-Cell RNA-seq Identifies Minority Cell Types of Kidney Distal Nephron. J. Am. Soc. Nephrol. 32, 886–896 (2021).

48. L. M. S. Gerhardt, A. P. McMahon, Multi-omic approaches to acute kidney injury and repair. Curr Opin Biomed Eng. 20 (2021), doi:10.1016/j.cobme.2021.100344.

49. R. Browaeys, W. Saelens, Y. Saeys, NicheNet: modeling intercellular communication by linking ligands to target genes. Nat. Methods. 17, 159–162 (2020).

50. M. Attanasio, N. H. Uhlenhaut, V. H. Sousa, J. F. O’Toole, E. Otto, K. Anlag, C. Klugmann, A.-C. Treier, J. Helou, J. A. Sayer, D. Seelow, G. Nürnberg, C. Becker, A. E. Chudley, P. Nürnberg, F. Hildebrandt, M. Treier, Loss of GLIS2 causes nephronophthisis in humans and mice by increased apoptosis and fibrosis. Nat. Genet. 39, 1018–1024 (2007).

51. S. Xu, X. Yang, Q. Chen, Z. Liu, Y. Chen, X. Yao, A. Xiao, J. Tian, L. Xie, M. Zhou, Z. Hu, F. Zhu, X. Xu, F. Hou, J. Nie, Leukemia inhibitory factor is a therapeutic target for renal interstitial fibrosis. EBioMedicine. 86, 104312 (2022).

52. S. Wang, X. Hu, L. Ma, L. Zhang, Y. Tian, CLCF1 is up-regulated in renal ischemia reperfusion injury and may associate with FOXO3. Ann Transl Med. 10, 399 (2022).

53. D. J. Kass, G. Yu, K. S. Loh, A. Savir, A. Borczuk, R. Kahloon, B. Juan-Guardela, G. Deiuliis, J. Tedrow, J. Choi, T. Richards, N. Kaminski, S. M. Greenberg, Cytokine-like factor 1 gene expression is enriched in idiopathic pulmonary fibrosis and drives the accumulation of CD4+ T cells in murine lungs: evidence for an antifibrotic role in bleomycin injury. Am. J. Pathol. 180, 1963–1978 (2012).

54. L. Crisponi, I. Buers, F. Rutsch, CRLF1 and CLCF1 in Development, Health and Disease. Int. J. Mol. Sci. 23 (2022), doi:10.3390/ijms23020992.

55. M. Murakami, D. Kamimura, T. Hirano, Pleiotropy and Specificity: Insights from the Interleukin 6 Family of Cytokines. Immunity. 50, 812–831 (2019).

56. A. Wehr, F. Tacke, The roles of CXCL16 and CXCR6 in liver inflammation and fibrosis. Curr. Pathobiol. Rep. 3, 283–290 (2015).

57. A. Wehr, C. Baeck, F. Heymann, P. M. Niemietz, L. Hammerich, C. Martin, H. W. Zimmermann, O. Pack, N. Gassler, K. Hittatiya, A. Ludwig, T. Luedde, C. Trautwein, F. Tacke, Chemokine receptor CXCR6-dependent hepatic NK T Cell accumulation promotes inflammation and liver fibrosis. J. Immunol. 190, 5226–5236 (2013).

58. M. Dudek, D. Pfister, S. Donakonda, P. Filpe, A. Schneider, M. Laschinger, D. Hartmann, N. Hüser, P. Meiser, F. Bayerl, D. Inverso, J. Wigger, M. Sebode, R. Öllinger, R. Rad, S. Hegenbarth, M. Anton, A. Guillot, A. Bowman, D. Heide, F. Müller, P. Ramadori, V. Leone, C. Garcia-Caceres, T. Gruber, G. Seifert, A. M. Kabat, J.-P. Mallm, S. Reider, M. Effenberger, S. Roth, A. T. Billeter, B. Müller-Stich, E. J. Pearce, F. Koch-Nolte, R. Käser, H. Tilg, R. Thimme, T. Boettler, F. Tacke, J.-F. Dufour, D. Haller, P. J. Murray, R. Heeren, D. Zehn, J. P. Böttcher, M. Heikenwälder, P. A. Knolle, Auto-aggressive CXCR6 CD8 T cells cause liver immune pathology in NASH. Nature. 592, 444– 449 (2021).

59. Y. Wu, C. An, X. Jin, Z. Hu, Y. Wang, Disruption of CXCR6 Ameliorates Kidney Inflammation and Fibrosis in Deoxycorticosterone Acetate/Salt Hypertension. Sci. Rep. 10, 133 (2020).

60. Z. Ma, X. Jin, L. He, Y. Wang, CXCL16 regulates renal injury and fibrosis in experimental renal artery stenosis. Am. J. Physiol. Heart Circ. Physiol. 311, H815–21 (2016).

61. M. Zhang, S. Zhang, T Cells in Fibrosis and Fibrotic Diseases. Front. Immunol. 11, 1142 (2020).

62. Y. Wang, J. Chang, B. Yao, A. Niu, E. Kelly, M. C. Breeggemann, S. L. Abboud Werner, R. C. Harris, M.-Z. Zhang, Proximal tubule-derived colony stimulating factor-1 mediates polarization of renal macrophages and dendritic cells, and recovery in acute kidney injury. Kidney Int. 88, 1274– 1282 (2015).

63. B. Venkatesan, A. Tumala, V. Subramanian, E. Vellaichamy, Transient silencing of Npr3 gene expression improved the circulatory levels of atrial natriuretic peptides and attenuated β-adrenoceptor activation-induced cardiac hypertrophic growth in experimental rats. Eur. J. Pharmacol. 782, 44–58 (2016).

64. M. Chromek, K. Tullus, O. Hertting, G. Jaremko, A. Khalil, Y.-H. Li, A. Brauner, Matrix metalloproteinase-9 and tissue inhibitor of metalloproteinases-1 in acute pyelonephritis and renal scarring. Pediatr. Res. 53, 698–705 (2003).

65. G. Kökény, Á. Németh, J. B. Kopp, W. Chen, A. J. Oler, A. Manzéger, L. Rosivall, M. M. Mózes, Susceptibility to kidney fibrosis in mice is associated with early growth response-2 protein and tissue inhibitor of metalloproteinase-1 expression. Kidney Int. 102, 337–354 (2022).

66. M. T. Rademaker, A. P. Pilbrow, L. J. Ellmers, S. C. Palmer, T. Davidson, P. Mbikou, N. J. A. Scott, E. Permina, C. J. Charles, Z. H. Endre, A. M. Richards, Acute Decompensated Heart Failure and the Kidney: Physiological, Histological and Transcriptomic Responses to Development and Recovery. J. Am. Heart Assoc. 10, e021312 (2021).

67. C. Song, S. Wang, Z. Fu, K. Chi, X. Geng, C. Liu, G. Cai, X. Chen, D. Wu, Q. Hong, IGFBP5 promotes diabetic kidney disease progression by enhancing PFKFB3-mediated endothelial glycolysis. Cell Death Dis. 13, 340 (2022).

68. M. Andreatta, J. Corria-Osorio, S. Müller, R. Cubas, G. Coukos, S. J. Carmona, Interpretation of T cell states from single-cell transcriptomics data using reference atlases. Nat. Commun. 12, 2965 (2021).

69. X. Wang, J. Chen, J. Xu, J. Xie, D. C. H. Harris, G. Zheng, The Role of Macrophages in Kidney Fibrosis. Front. Physiol. 12, 705838 (2021).

70. P. E. Cippà, J. Liu, B. Sun, S. Kumar, M. Naesens, A. P. McMahon, A late B lymphocyte action in dysfunctional tissue repair following kidney injury and transplantation. Nat. Commun. 10, 1157 (2019).

71. M. Adler, A. Mayo, X. Zhou, R. A. Franklin, M. L. Meizlish, R. Medzhitov, S. M. Kallenberger, U. Alon, Principles of Cell Circuits for Tissue Repair and Fibrosis. iScience. 23, 100841 (2020).

72. S. Shah, Y. Takei, W. Zhou, E. Lubeck, J. Yun, C.-H. L. Eng, N. Koulena, C. Cronin, C. Karp, E. J. Liaw, M. Amin, L. Cai, Dynamics and Spatial Genomics of the Nascent Transcriptome by Intron seqFISH. Cell. 174, 363–376.e16 (2018).

73. C.-H. L. Eng, S. Shah, J. Thomassie, L. Cai, Profiling the transcriptome with RNA SPOTs. Nat. Methods. 14, 1153–1155 (2017).

74. Y. Takei, J. Yun, S. Zheng, N. Ollikainen, N. Pierson, J. White, S. Shah, J. Thomassie, S. Suo, C.-H. L. Eng, M. Guttman, G.-C. Yuan, L. Cai, Integrated spatial genomics reveals global architecture of single nuclei. Nature. 590, 344–350 (2021).

75. Y. Hao, S. Hao, E. Andersen-Nissen, W.M. Mauck 3rd, S. Zheng, A. Butler, M. J. Lee, A. J. Wilk, C. Darby, M. Zager, P. Hoffman, M. Stoeckius, E. Papalexi, E. P. Mimitou, J. Jain, A. Srivastava, T. Stuart, L. M. Fleming, B. Yeung, A. J. Rogers, J. M. McElrath, C. A. Blish, R. Gottardo, P. Smibert, R. Satija, Integrated analysis of multimodal single-cell data. Cell. 184, 3573–3587.e29 (2021).

76. I. Korsunsky, N. Millard, J. Fan, K. Slowikowski, F. Zhang, K. Wei, Y. Baglaenko, M. Brenner, P.-R. Loh, S. Raychaudhuri, Fast, sensitive and accurate integration of single-cell data with Harmony. Nat. Methods. 16, 1289–1296 (2019).

